# Phase-precession-like Effect in the Anterior Insula Cortex during Reward Expectancy

**DOI:** 10.1101/2023.09.19.558205

**Authors:** Linglin Yang, Katia Lehongre, Hongyi Ye, XinFeng Yu, Vincent Navarro, Sen Cheng, Nikolai Axmacher, Shuang Wang, Hui Zhang

## Abstract

Reward expectancy shapes behavioral performance by coordinating the allocation of cognitive resources, which incorporates the involvement of the anterior insular cortex (AIC). To investigate the electrophysiological mechanisms of the AIC during reward expectancy, we collected intracranial electroencephalographic data from epilepsy patients undergoing clinical monitoring. Subjects navigated a virtual T-maze containing rewards at three predetermined locations. We focused on the period immediately before entering the reward zone, termed the reward expectancy stage. During this stage, we identified the reward-specific brain patterns (RBPs) that were preactivated in the AIC. This pre-activation preceded the emergence of robust phase-amplitude coupling (PAC), where theta oscillations (5-7 Hz) modulated gamma activities (75–110 Hz). The PAC strength correlated positively with the pre-activation level of RBPs, implying the potential role of oscillatory coordination in amplifying preactivated reward representations. Strikingly, this PAC effect exhibited a specific temporal structure that peak gamma activity became progressively coupled to earlier theta phases as reward approached, mirroring the theta phase precession phenomenon previously observed in frontal and temporal lobes. We refer to this dynamic as phase-precession-like effect (PPLE). Moreover, subjects exhibiting PPLE in the AIC presented greater trial-by-trial improvements in response latency. Taken together, these findings shed light on the electrophysiological mechanisms of the AIC underlying reward expectancy.

## Introduction

Many daily behaviors of humans are frequently driven by the expectancy of rewards, whether material or spiritual^1,2^. Reward expectancy has been proposed to boost behavioral performance^2^, particularly when the expected reward is ultimately obtained^3,4^. This process hinges on context-reward associations, wherein rewards become linked to specific environmental or sensory cues^5^. Recognition of a reward-associated context triggers activation of the neural pattern encoding reward memory^3^, prompting the brain to allocate heightened attentional and motivational resources to detect expected rewards within the reward-associated contexts ^2–4,6^. Subsequent reward delivery consistent with expectations reinforces these associations via feedback-dependent plasticity^3,7,8^. Critically, reward expectancy integrates salience detection, cognitive resource allocation, and predictive coding - processes coordinated by the salience network (SN)^3,9,10^. As a critical component of the SN, the anterior insula cortex (AIC) activates ahead of other cortical areas during various cognitive tasks, such as the oddball attention task and flanker task^11,12^. These preceding neural activities in the AIC are speculated to participate in integrating multimodal information and prioritizing cognitive resources for reward expectancy^13–15^. Electrophysiological recordings in non-human primates have revealed neuronal activities related to reward expectancy in the AIC, showing that approximately a quarter of responsive neurons are activated preceding reward delivery^16^. These findings have been extended to humans in a more recent study, demonstrating AIC engagement during reward expectancy^10^. Nevertheless, the precise electrophysiological mechanism by which the AIC mediates reward expectancy remain poorly characterized. Elucidating these mechanisms - particularly the temporal coordination of neural activity during anticipation - could advance strategies to optimize information processing speed and cognitive efficiency in goal-directed behaviors. Intracranial electroencephalography (iEEG) captures high-temporal-resolution neural activity that unveils the rapid dynamics during cognitive processes in deep brain regions like the AIC. A hallmark of such dynamics is phase-amplitude coupling (PAC), where high-frequency activities are synchronized to specific phases of low-frequency oscillations (LFOs), especially theta oscillations^17–19^. Theta rhythms facilitate inter-regional communication information by entraining local gamma generators across distributed networks^20–22^, while enhanced PAC effect promotes information integration^19,23^, memory consolidation^18^, and context-item binding^17,24^.

Basic characteristics of PAC may reflect the functional configuration of a neural network during a given task. The timing of high-frequency bursts relative to LFO phases carries distinct neural signatures implicated in a range of cognitive tasks^25,26^. For instance, gamma activities coupled to the peak of theta oscillations contribute to memory encoding, while those coupled to the trough of theta oscillations participate in memory retrieval^27–29^. During sequential event encoding, distinct LFO phases in frontal and temporal lobes track different events showing that the neural pattern underlying each event is associated with different phases of LFOs^18,21,30–35^. Such a phase-locking phenomenon is assumed to be the neural foundation of the limited capacity of short-term memory^18,21,36,37^. Notably, applying transcranial alternating current stimulation (tACS) over the sensorimotor cortex to induce pronounced gamma activities at the peak of theta oscillations can substantially enhance motor skill acquisition^38^. These findings collectively suggest that the PAC-mediated modulation reflects the neural signatures of brain regions underlying specific cognitive processes.

Moreover, another temporal organizing principle – phase precession – has been observed in rodents and humans. This phenomenon characterizes as a progressive shift in spike timing to earlier theta phases as subjects traverse place fields^39–41^. Initially identified in the hippocampus and entorhinal cortex, phase precession has since been documented in reward-related regions (e.g., ventral striatum, lateral septum), where anticipatory activity toward reward sites correlates with phase precession^42–45^. In humans, non-spatial phase precession has been revealed in the temporal and frontal regions during goal-directed tasks, hinting its broader role in predicting coding^41,46^.

Existing evidence on phase precession derives predominantly from microscale single-unit recordings. It remains unclear whether and how this phenomenon manifests at macroscale neural levels (e.g., population-level oscillations) or interfaces with reward anticipation. To address this gap, we analyzed iEEG data from presurgical epilepsy patients navigating a virtual T-maze with pre-defined rewards. By integrating representational similarity analysis (RSA) and PAC analysis, we investigated the neural dynamic in the AIC during the period preceding reward delivery. Our study aims to connect macroscale oscillatory dynamics with microscale phase precession phenomena, advancing understanding of how neural systems anticipate and optimize behavior for reward acquisition.

## Results

To investigate the electrophysiological mechanisms of reward expectancy in the AIC, we tested nine epilepsy patients with 25 AIC contacts of implanted electrodes (18 in the right hemisphere; Figure 1A and Table S1). As a control, we also examined 19 hippocampal contacts (9 in the right hemisphere), which included 7 contacts from 3 patients with AIC contacts and the remaining 12 contacts from 4 new patients. AIC and HPC contacts demonstrated comparable lateralization distributions (χ^2^*_(1, 44)_* = 2.76, *p* = 0.12). Subjects were asked to navigate a virtual T-maze and collect rewards while iEEG activities were recorded (Figure 1B-D). Subsequently, we examined iEEG activities aligned to three key events: 1) entering the reward zone (ERZ, experimental condition - reward appearance on the monitor Figure 1D), 2) picking up the reward (PR, control condition), and 3) entering the decision zone (EDZ, control condition).

**Figure 1.**
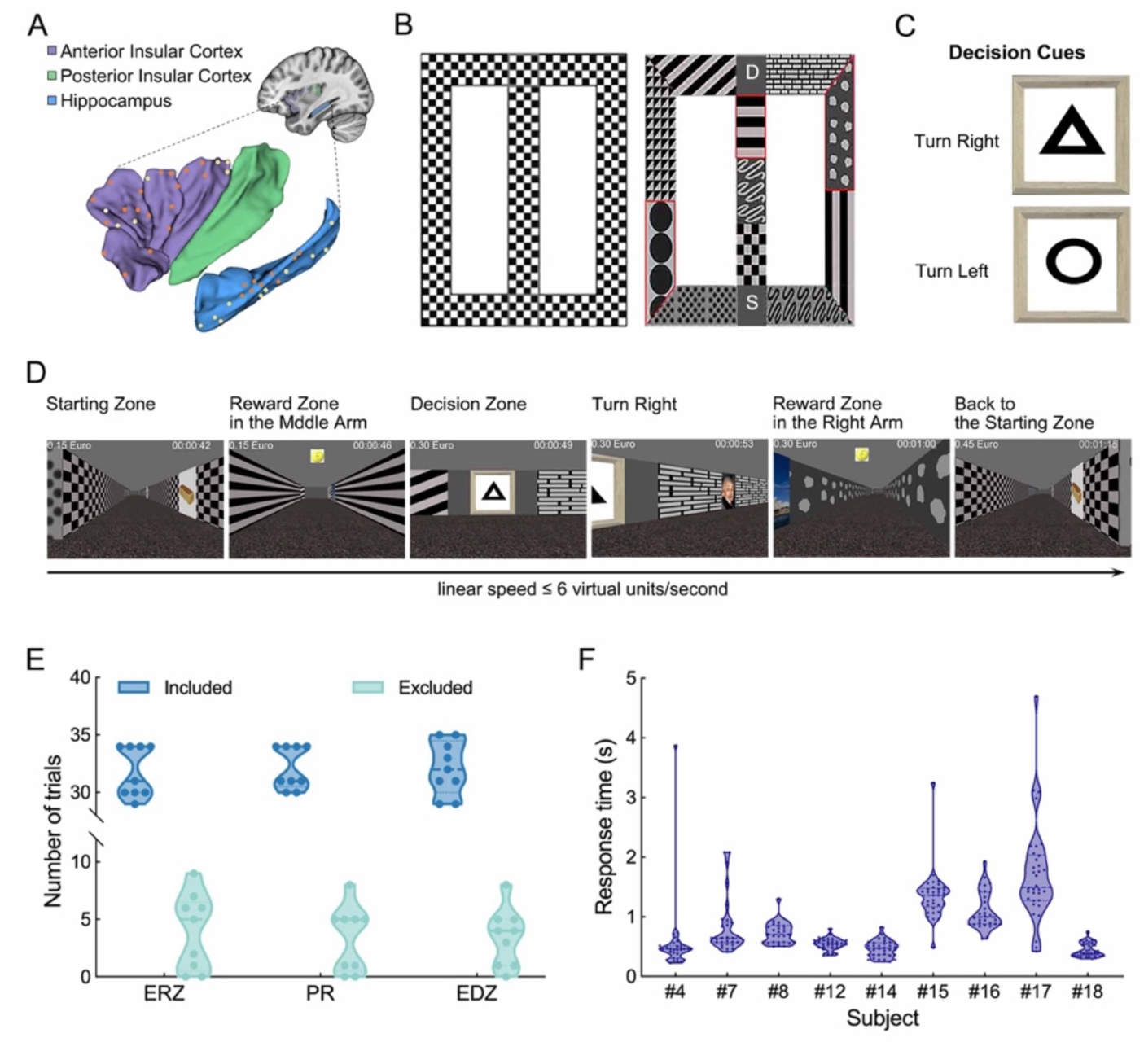
Contact locations, experimental paradigm, and behavioral performance. (A) Spatial distribution of contacts in the anterior insular cortex (AIC, purple), posterior insular cortex (PIC, green), and hippocampus (HPC, blue). For illustration purposes, contacts from both hemispheres are drawn together. Each orange sphere indicates a contact in the right hemisphere, and each yellow sphere indicates a contact in the left hemisphere. (B) Overhead view of the T maze. Left panel: Diagram of the virtual maze. Each small square represents one square virtual unit. Right panel: T maze adorned with various patterns on its walls. The gray square marked with “S” represents the starting zone where each trial starts and ends, while the gray square marked with “D” represents the decision zone where subjects are expected to choose the correct direction according to the cue. Areas circled with red lines represent reward zones, where subjects are expected to collect gold coins by pressing the Enter button as soon as possible. Otherwise, the reward disappears when they are outside of the reward zone. (C) Cues on the wall of the decision zone. Circle represents turning left and triangle represents turning right. (D) An example trial from the first-person perspective in the maze. (E) Included (blue) and excluded (green) trials are similar across conditions. Note that there is a discontinuity in the y-axis to enhance visualization. ERZ, entering the reward zone; PZ, picking up the reward; EDZ, entering the decision zone. Each dot within violin plots indicates a subject. (F) Behavioral performance for subjects with AIC contacts. Each violin plot indicates one subject, and each dot within violin plots indicates one trial.

This hierarchical design enabled targeted analysis of reward-processing mechanisms while controlling for motor execution (PR) and baseline spatial navigation (EDZ). The number of included trials for different conditions were matched (31.78 ± 2.17 [mean ± SD] for ERZ, 32.11 ± 1.83 for PR, and 31.89 ± 2.32 for EDZ; *F_(2, 24)_* = 0.058, *p* = 0.94, one-way ANOVA; Figure 1E and Table S1; see *Star Method - IEEG Data Acquisition and Preprocessing*). For each trial, subjects’ response time (RT) to pick-up rewards were calculated for further analyses (Figure 1F; see *Results - Functional Relevance of PPLE*).

### Pre-activation of the Reward-specific Brain Patterns when Approaching Rewards

Reward expectancy indicates that the brain starts to process reward-relevant information before the onset of rewards. Thus, we hypothesized that the brain patterns representing rewards would be preactivated prior to actual reward delivery. To test this hypothesise, we investigated the pre-activation of reward-specific brain patterns (RBPs) in AIC and hippocampal contacts before reward onset using representational similarity analysis (RSA)^47–50^. To identify RBPs, we first computed the neural pattern similarities during reward viewing (post-ERZ periods) across different spatial locations (the central and two lateral roads; RSA_ERZ/ERZ_) to control for location-specific neural representations (Figure 1B, right panel; see *Star Methods - Representational Similarity Analysis and Pre-activation of Reward-Specific Brain Patterns*). As a control comparison, we then calculated the pattern similarities between reward viewing (post-ERZ periods) and non-reward viewing periods (post-EDZ periods; RSA_ERZ/EDZ_; Figure 1B-D and 2A). Finally, we derived RBPs by contrasting these similarity measures (RSA_ERZ/ERZ_ vs. RSA_ERZ/EDZ_) during post-ERZ periods (Figure 2B), ensuring the identified patterns specifically reflected reward processing rather than spatial or task-general neural activities. RBPs exhibited distinct temporal dynamics across regions during reward processing. In the AIC, neural activity within 0.875–1.3 seconds following reward onset showed significantly higher neural pattern similarity for reward viewing pairs (RSA_ERZ/ERZ_ values) compared to reward viewing and non-reward viewing pairs (RSA_ERZ/EDZ_ values, *p_corrected_* = 0.047; Figure 2B, top panel). In contrast, the hippocampus encoded reward information earlier, with enhanced similarity for reward viewing (RSA_ERZ/ERZ_ values) versus reward viewing and non-reward viewing pairs (RSA_ERZ/EDZ_ values) emerging within 0.1–0.525 seconds post-reward onset (*p_corrected_* = 0.033; Figure 2B, bottom panel).

**Figure 2.**
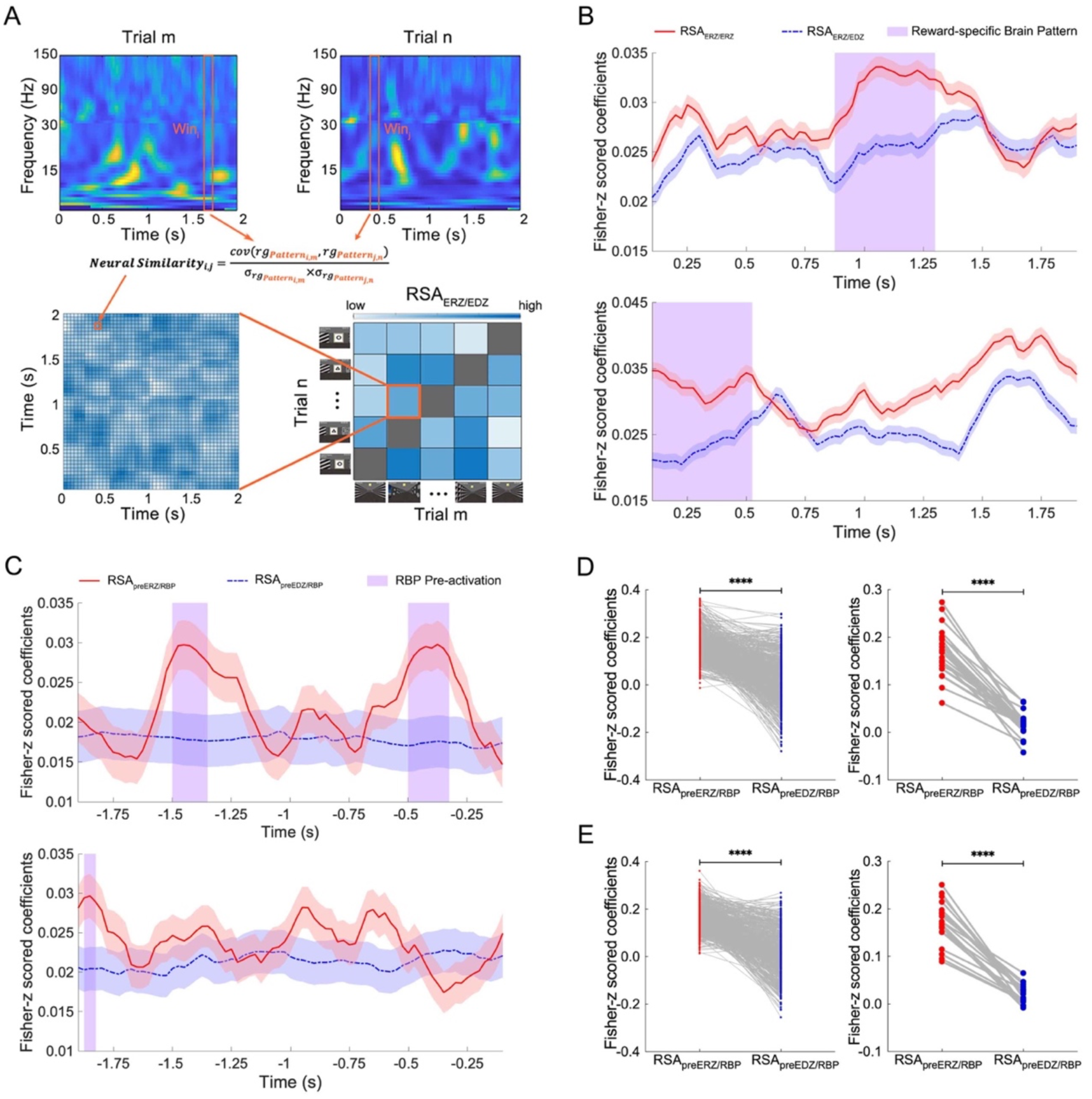
Pre-activation of reward-specific neural patterns before entering reward zones in the AIC and hippocampus. (A) Scheme of representational similarity analysis (RSA) for reward-specific brain patterns (RBPs). Representational similarity matrix (bottom left) illustrates the neural similarities of the spectrum within each time window between different trials (top). Representational similarity matrix of RSA_ERZ/EDZ_ (bottom right) displays the categorical information structure of different stimuli (rewards or geometric figures). ERZ, entering the reward zone; EDZ, entering the decision zone. (B) Time-resolved category specificity revels RBPs in the anterior insular cortex (AIC, top) and hippocampus (bottom). Each line represents time courses of neural similarity (mean ± SEM) for ERZ/ ERZ (red) and ERZ/ EDZ (blue) condition pairs. Purple shading indicates time windows carrying RBPs (i.e., RSA_ERZ/ERZ_ > RSA_ERZ/EDZ_). X-axis denotes time course relative to the post-ERZ epoch. (C) Pre-activation of RBPs before entering reward zones in the AIC (top panel) and hippocampus (bottom panel). Each line represents time-resolved similarity trajectories (mean ± SEM) for preERZ/RBP (red) and preEDZ/RBP (blue) comparisons. Purple shading highlights pre-ERZ time windows for RBP pre-activation (i.e., RSA_preERZ/RBP_ > RSA_preERZ/RBP_). X axis indicates the time course relative to pre-ERZ epoch. (D and E) Differential pre-activation of RBPs between preERZ/RBP and preEDZ/RBP in the AIC (D) and hippocampus (E). Left panels illustrate trial-level differences, with red spheres representing RSA_preERZ/RBP_ values for individual trials and blue spheres denoting RSA_preEDZ/RBP_ values. Right panels depict contact level analyses, where red spheres denote trial-averaged RSA_preERZ/RBP_ values at individual recording contact; blue spheres represent trial-averaged RSA_preEDZ/RBP_ values. ****, *p*<0.001.

To examine the pre-activation of RBP, we computed the neural similarities between each time window before reward onset (pre-ERZ periods) and each time window showing RBP when viewing rewards (post-ERZ periods; RSA_preERZ/RBP_). To control for location-specific effects, we compared brain pattern from pre-ERZ periods at the central road and post-ERZ periods at the lateral roads. To avoid the temporal proximity effect (i.e., autocorrelations), correlations between pre- and post-ERZ periods from the same trial were excluded. As a control measure, we calculated the neural similarities between the time windows before decision sign onset (pre-EDZ periods) and time windows showing RBP when viewing rewards (post-ERZ periods; RSA_preEDZ/RBP_). The results revealed distinct temporal profiles of RBPs pre-activation between the AIC and hippocampus (Figure 2C), as described above. Strikingly, temporally refined analysis revealed two pre-ERZ time windows of RBP pre-activation in the AIC (one from 1.35 to 1.5 seconds and another from 0.325 to 0.5 seconds before the onset of reward, *ps_corrected_* = 0.001), where the RSA_preERZ/RBP_ values were higher than the average RSA_preEDZ/RBP_ values (Figure 2C, top panel). The hippocampus, however, exhibited even earlier anticipatory dynamics, with RBP pre-activation peaking from 1.825 to 1.875 seconds before the onset of reward (*p_corrected_* = 0.001; Figure 2C, bottom panel). To quantify the pre-activation of RBPs, we extracted the maximum neural pattern similarity values (RSA_preERZ/RBP_ and RSA_preEDZ/RBP_) within each trial, separately for the AIC and hippocampus. In the AIC, the pre-activation levels of RBPs during pre-ERZ periods (0.17 ± 2.15×10^-3^, mean ± SEM) significantly exceeded those during pre-ERZ periods (0.02 ± 3.65×10^-3^) at the trial level (*t_(1526)_* = 36.33, *p* < 0.0001, two-sample t test; Figure 2D, left). A parallel effect was observed in the hippocampus (RSA_preERZ/RBP_: 0.17 ± 2.30×10^-3^ vs. RSA_preEDZ/RBP_: 0.02 ± 3.80×10^-3^, *t_(1264)_* = 33.54, *p* < 0.0001; Figure 2E, left). These result were validated by collapsing trails within individual contacts (AIC: 0.17 ± 9.62×10^-3^ vs. 0.02 ± 4.63×10^-3^, *t_(24)_* = 15.94, *p* < 0.0001; hippocampus: 0.17 ± 0.01 vs. 0.02 ± 4.39×10^-3^, *t_(18)_* = 13.81, *p* < 0.0001; paired-t test; Figure 2D and E, right panels).

Together, these results showed that existence of RBPs in both the AIC and hippocampus after the onset of rewards. Critically, these RBPs were robustly pre-activated prior to the onset of rewards, with distinct temporal dynamics between regions. This spatiotemporally precise pre-activation underscores conserved reward-predictive mechanisms across neural scales. Strikingly, the RBP pre-activation in the AIC suggests that it started to process reward information before the appearance of reward, implicating it in reward expectancy.

### Coupling between Theta Phases and Gamma Amplitudes in the AIC when Approaching Rewards

To estimate the neural dynamics of the AIC during reward expectancy, we compared the PAC effect before and after entering the reward zone (pre- and post-ERZ events). Firstly, the time course of phases for each low-frequency (frequency_low_;1-12Hz in 1 Hz steps with a 1Hz-bandwidth; see *Supplementary Information-Spectral Dynamics of Theta and Alpha Oscillations in the AIC and Hippocampus*; Figure S1) and the time course of amplitudes for each high-frequency (frequency_high_; 30-150Hz in 5 Hz steps with a 24Hz-bandwidth) were extracted for both the time windows of pre- and post-ERZ events for each trial. Utilizing a method that has been previously described^51^, PAC strength was subsequently calculated between each low-frequency band and each high-frequency band for both the pre- and post-ERZ events within each trial (see *Star Methods - Phase-Amplitude Coupling Analysis and Identification of Peak Gamma Activities*). Using a multilevel linear regression analysis by considering the variance at contact and subject levels, we computed the difference of PAC values between pre- and post-ERZ events for each frequency_low_-frequency_high_ pair. To correct errors caused by multiple comparisons, we used a cluster-based permutation analysis to extract frequency_low_-frequency_high_ clusters showing significant coupling when subjects were approaching rewards^52^. We found that the coupling between theta phases (5-7 Hz) and gamma amplitudes (75-110 Hz) was stronger during pre-ERZ events compared to post-ERZ events (*p_corrected_* = 0.005, Figure 3A). We applied the same analysis to control conditions (i.e., PR and EDZ events) in AIC contacts and to all conditions in hippocampal contacts but did not find any significant clusters (Figure S2A). These results suggest that PAC effect occurs when approaching rewards, specifically in the AIC.

**Figure 3.**
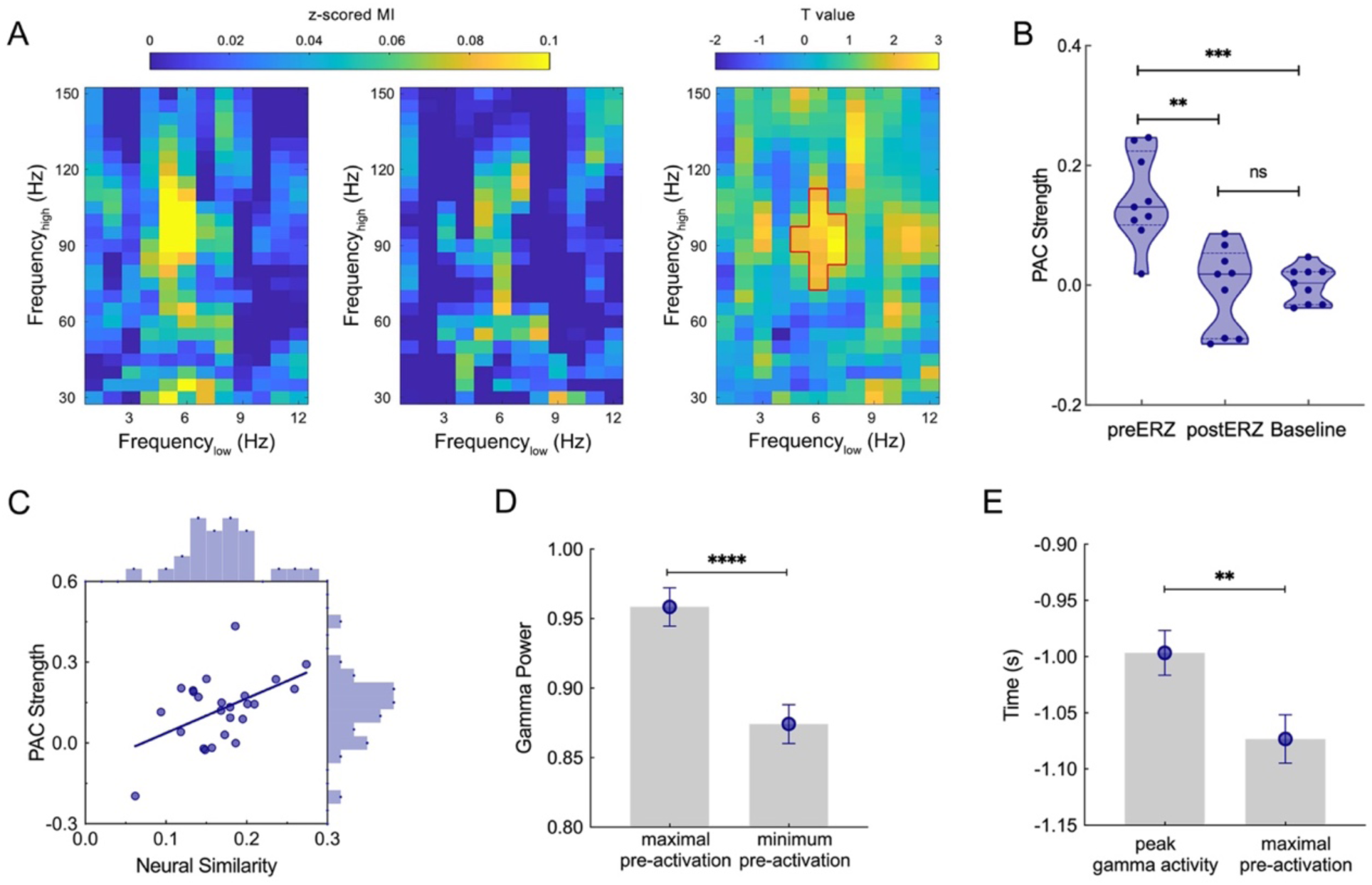
Phase-amplitude coupling (PAC) and its relationship to RBP pre-activation in the anterior insular cortex (AIC) before entering reward zones. (A) Comodulograms illustrate the z-scored modulation index (MI) corresponding to the pre-ERZ (left) and post-ERZ (middle) events. The cluster encircled in red lines (right) highlights a more pronounced PAC effect in pre-ERZ events as compared to post-ERZ events, (frequency_low_: 5-7Hz; frequency_high_: 75-110Hz). ERZ, entering the reward zone. (B) Average PAC values within the frequency_low_-frequency_high_ cluster from panel A for individual subjects in the pre-ERZ, post-ERZ, and baseline conditions. Each violin plot indicates one condition, and each dot within violin plots indicates one subject. **, 0.005≤*p*<0.01. (C) Neural similarities of pre-ERZ event relative to reward-specific brain patterns (RBP) positively correlate with the strength of PAC. Each sphere indicates one contact, blue line indicates the fitted line. (D) Gamma power showing PAC effects is higher at the time bins showing maximal pre-activation of RBP than those showing minimum pre-activation. Error bar indicates SEM. ****, *p*<0.001. (E) Peak gamma activities showing PAC effects occur later than RBP pre-activation of. Y axis indicates the time course relative to pre-ERZ epoch. Error bar indicates SEM. **, 0.005≤*p*<0.01.

It is plausible that the observed enhanced coupling between theta phases and gamma amplitudes within the AIC during pre-ERZ periods contributes to reward expectancy. However, the enhanced PAC effect observed may potentially be attributed to a reduction in PAC strength during the post-ERZ periods. To address this possibility and ensure the robustness of our findings, we extracted the mean PAC values within the significant frequency_low_-frequency_high_ cluster identified earlier, for the pre-ERZ period, post-ERZ period and baseline which was randomly selected time periods outside ERZ periods, respectively. As expected, the PAC strength during the pre-ERZ period (0.14 ± 0.07, [mean ± SD]) was significantly greater than that observed during the post-ERZ period (−6×10^-3^ ± 0.07, *t_(8)_* = 3.73, *p* = 5.8×10^-3^, paired-t test) and at baseline (10^-3^ ± 0.03, *t_(8)_* = 6.42, *p* = 2×10^-4^; Figure 3B). No significant difference in PAC strength was found between the post-ERZ period and baseline (*t_(8)_* = 0.24, *p* = 0.81). These results suggest that the enhanced PAC effect in the AIC is not merely a consequence of reduced PAC strength in the post-ERZ period, but rather reflects a genuine increase in coupling strength during the pre-ERZ period. Taken together, these results suggested that gamma band activities coupled to theta oscillations during pre-ERZ period within each trial, specifically for contacts located in the AIC.

### PAC Effects were Related to Pre-activation of Reward-specific Brain Patterns

As reported above, both RBP pre-activation and theta-gamma coupling occurred when approaching rewards. Thus, we investigated the relationship between them. The average PAC value and the mean neural similarity extracted from individual trials were found to be positively correlated (*r* = 0.49, *p* = 0.012, Pearson’s correlation; Figure 3C). We subsequently compared the gamma power between time bins exhibiting the maximal and minimum pre-activation levels of RBP within each trial. Gamma power was significantly elevated during epochs of maximal RBP pre-activation compared to trough epochs (maximal: 0.96 ± 0.01 [mean ± SEM], minimum: 0.87 ± 0.01, *p* < 10^-4^, Wilcoxon matched-pairs signed rank test; Figure 3D). These findings indicated that RBP pre-activation and PAC effect were correlated in strength.

Furthermore, we investigated the temporal relationship between the peak gamma power derived from PAC analysis and RBP pre-activation. The peak gamma power relative to the preferred theta phase occurred at 1.00 ± 0.02 second before the onset of rewards, markedly later than the highest pre-activation (1.07 ± 0.02 second before the onset of rewards; *p* = 5.4×10^-3^, Wilcoxon matched-pairs signed rank test; Figure 3E). This temporal dissociation indicates that RBP pre-activation precedes gamma-PAC dynamics. Taken together, these findings indicate that the strength of PAC is positively correlated with the level of RBP pre-activation, indicating the potential relationship between these two reward-expectancy-related effects. Critically, RBP pre-activation occurs prior to the peak PAC effect, suggesting that RBPs functionally anticipates the emergence of theta-gamma coordinated activity during reward expectancy.

### Gamma Activities Exhibit Phase-precession-like Effects in the AIC when Approaching Rewards

The result reported above suggests that gamma activities are coupled to theta oscillations across trials when subjects are approaching the same rewards. Notably, the theta phases coupled to gamma activity demonstrated trial-to-trial variability within individual recording contacts (Figure 4A and S4A), reflecting dynamic reconfiguration of phase-amplitude coordination across behavioral epochs. Hereby we looked further into the characteristic of PAC data to examine the dynamics of the maximal theta phases across trials. We therefore aimed to investigate if gamma activities were coupled to theta phases that systematically shifted forward or backward when subjects were approaching a reward. To this end, we performed a circular-linear regression analysis between maximal theta phases of PAC effects during the pre-ERZ periods and the number of trials in individual AIC contacts (See *Star Methods - Phase-Precession-Like Effect of Peak Gamma Activities*)^41^. If the peak gamma power is coupled to gradually shifted theta phases when approaching rewards, we would expect a positive or negative correlation between the maximal theta phase (circular data) and the trial number (linear data; Figure 4A). Indeed, we observed a significant negative correlation in 9 out of 25 (36%) AIC contacts from 6 out of 9 (66.7%) subjects and no correlations in the other 16 contacts (Figure 4B-D and S4A). The prevalence of such a negative correlation at both contact and subject levels were significantly higher than chance level (i.e., 5%; *ps* ≤ 10^-6^, Binomial test)^41^. It suggests that peak gamma activities are coupled to gradually earlier theta phases when approaching the same reward. The phenomenon is reminiscent of the phase prerecession effect and termed as phase-precession-like effect (PPLE).

**Figure 4.**
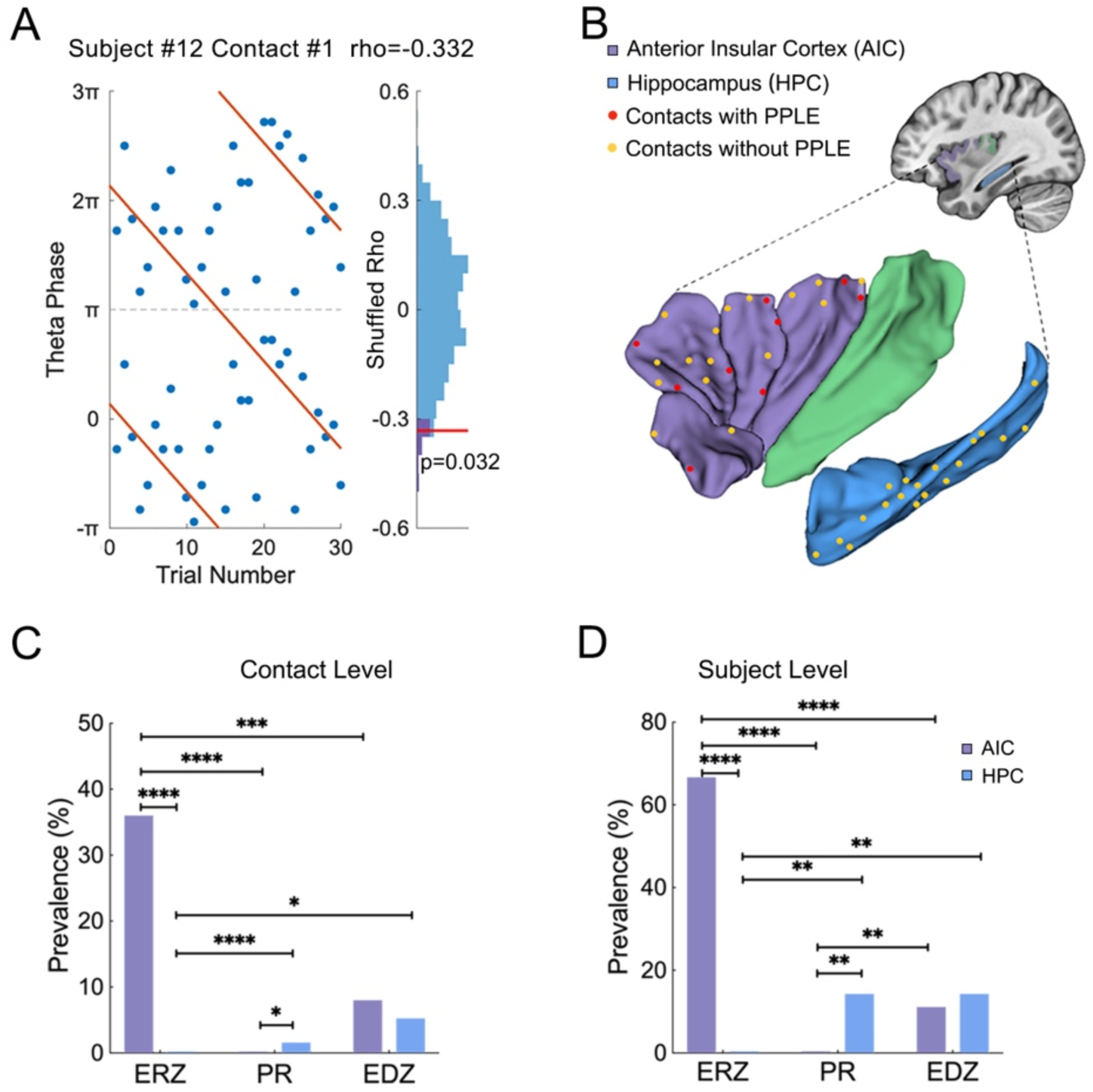
Phase-precession-like effect (PPLE) in the anterior insular cortex (AIC) before entering reward zones. (A) An example demonstrates that peak gamma activities are locked to earlier theta phases over trials when subjects are approaching the same reward zone. Blue spheres indicate individual trials, and orange lines indicate the fitted line between theta phases corresponding to peak gamma activities and trial numbers. Note that theta phases are duplicated vertically to enable visualization of circular-linear regression. The statistical assessment of circular-linear regression rho is conducted using surrogate distribution of circular-linear regression rho values generated by permutation of trial labels. Red lines indicate values of real data, and purple shading indicate the 95^th^ percentile of the surrogate distribution. Spatial distribution of contacts showing PPLE in the AIC and hippocampus. (C and D) Prevalence of PPLE in the AIC (purple) and hippocampus (blue) for each condition at contact (C) and subject (D) levels. ERZ, entering the reward zone; PR, picking up the reward; EDZ, entering the decision zones. *, 0.01≤*p*<0.05; **, 0.005≤*p*<0.01; ***, 0.001≤*p*<0.005; ****, *p*<0.001.

It is possible that PPLE was caused by systermatical shift of peak gamma activities across trials. To test this possiblity, we performed simple linear regression to assess the relationship between the timepoints corresponding to peak gamma activities and trial sequence (Figure S5). If gamma peaks shifted progressively earlier over trials, a negative correlation would emerge between gamma peak timing and trial number (hypothesis: *β* < 0). After shuffling trial labels 500 times, no AIC contacts exhibited significant negative correlations (*Ωs* ≥ −0.35, *ps_corrected_* ≥ 0.064), while one recording contact (from subject #16) exhibited a positive correlation (*β* = 0.40, *p_corrected_* = 0.036). It argues against systematic temporal forward shift in gamma activity as the driver of PPLE, suggesting that PPLE primarily reflects an internal neural activity associated with reward expectancy, rather than being driven by the estimation of reward onset timing.

The PPLE in the AIC when approaching rewards was possibly the electrophysiological foundation of reward expectancy. Meanwhile, it might also relate to the expectancy of visual stimuli, specific locations, or outcomes of picking up rewards. To determine whether PPLE in the AIC was specifically related to reward expectancy, we compared the prevalence of PPLE during pre-ERZ events with two control conditions (i.e., pre-PR or pre-EDZ events; Figure 4C-D, S6A and S7A). PPLE occurred in fewer AIC contacts during pre-PR events (0/25) and pre-EDZ events (2/25) which were significantly smaller than during pre-ERZ events (pre-PR: *p* = 1.4×10^-5^; pre-EDZ: *p* = 0.0016; Binomial test). At the subject levels, a similar pattern was observed, demonstrating a lower occurrence of PPLE during pre-PR events (0/9 cases, *p* = 4.6×10^-5^) and pre-EDZ events (1/9 cases, *p* = 8.9×10^-4^) in comparison to pre-ERZ events. These findings suggest that PPLE in the AIC is preferred to reward expectancy rather than reward outcomes (i.e., pre-PR events) or non-reward visual stimuli or specific locations (i.e., pre-EDZ events). To determine if PPLE prior to reward appearance is specific to the AIC, we examined the prevalence of PPLE in hippocampal contacts during the pre-ERZ event (Figure 4B-D and S4B). None of the hippocampal contacts (0/19) exhibited PPLE, showing significantly lower prevalence compared to the AIC contacts (*p* = 2.1×10^-4^, Binomial test). This dissociation maintained statistical significance at the subject level, with no subjects (0/7) exhibiting PPLE in hippocampal contacts compared to those with PPLE in AIC contacts (6/9, *p* = 4.3×10^-4^). It suggests that PPLE during pre-ERZ periods is more prevalent in the AIC rather than in the hippocampus.

All together, the PPLE reveals that gamma activities peak at progressively earlier theta phases when subjects approaching rewards. Futhermore, such a reward-expectancy-related PPLE is especially prominent in the AIC.

### Functional Relevance of PPLE

The high prevalence of PPLE in the AIC preceding ERZ events suggests a potential role for PPLE in reward expectancy. Previous studies have shown that reward anticipation facilitates cognitive performance, such as shortening response latency^6,53^. We thus investigated if PPLE facilitated subjects’ responses to rewards. In order to examine the improvement of RT across trials, z-transformed RT was extracted for each trial in each subject (Figure S8). Trials with a z-score > 3 were excluded from further data analysis. We then performed a linear regression analysis to predict the z-transformed RT of picking up rewards using the number of trials. RTs trended down for all nine subjects with AIC contacts as the number of trials increased (*βs* < 0; Figure 5A and S8). We tested the significance of RT decline pattern by performing a surrogate analysis on each subject. Within each permutation, we randomly shuffled the trial numbers, computed the linear regression again, and obtained a surrogate coefficient. The permutation procedure was repeated 500 times, generating 500 surrogate coefficients. We then ranked the empirical coefficient within the distribution of surrogate coefficients for each subject. We observed a significant decrease in RT with increasing number of trials in six subjects (*ps* < 0.05; Figure S8).

**Figure 5.**
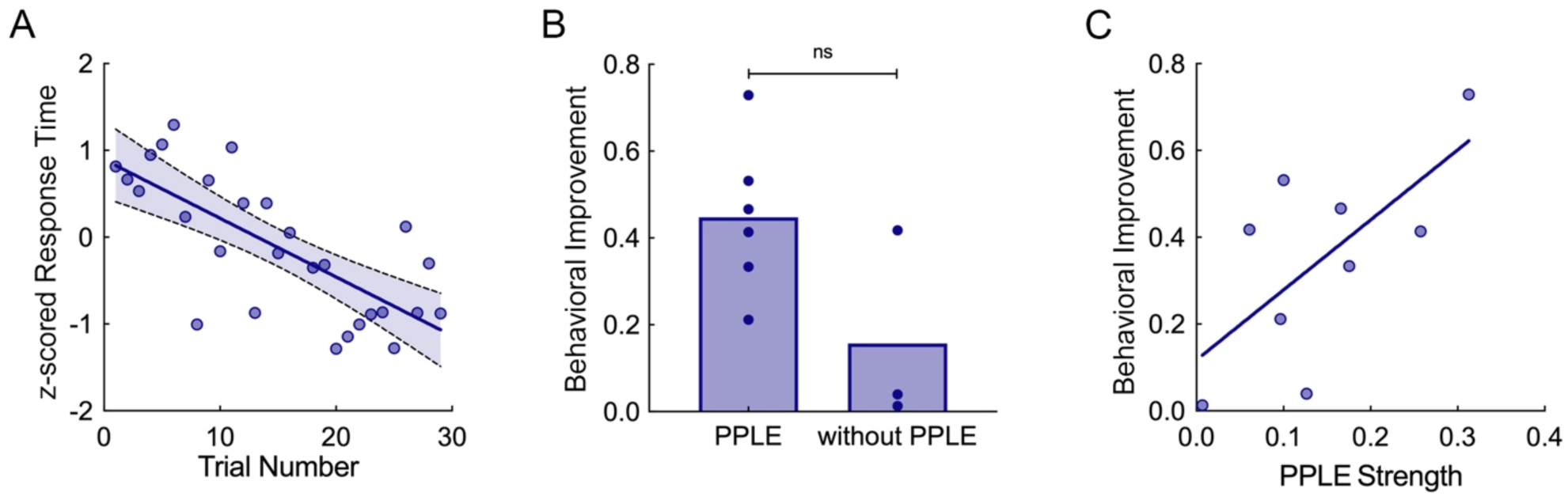
Functional relevance of phase-precession-like effect (PPLE). (A) An example showing that response time (RT) decreases with increased trial numbers. Each sphere indicates one trial, blue line indicates the fitted line between z-scored RT and trial numbers, and shaded area represents 95% confidence interval of the fitted line. (B) No significant difference in behavioral improvement is found between subjects with PPLE and those without PPLE. Each sphere indicates one subject. (C) Behavioral performance improved greater for subjects with stronger PPLE. Each sphere indicates one subject, and blue line indicates the fitted line between behavioral improvement and PPLE strength.

To investigate the potential relationship between PPLE and behavioral performance, we extracted the strength of PPLE and the improvement in behavioral performance for further analysis. PPLE strength for each subject was determined by calculating the average absolute value of the circular-linear coefficients across AIC contacts for PPLE. The parameter quantifying behavioral improvement was defined as the absolute regression coefficient derived from a linear model predicting z-transformed RTs using trial numbers (Figure S8). Firstly, we compared behavioral improvement between subjects with and without PPLE. Behavioral improvement did not differ significantly between subjects with PPLE (0.45 ± 0.18 [mean ± SD]) and those without PPLE_PAC_ (0.16 ± 0 .23, *p* = 0.17, Wilcoxon matched-pairs signed rank test; Figure 5B). Additionally, we conducted correlation analyses between the strength of PPLE and the magnitude of performance improvement across subjects. A significant positive correlation was observed between PPLE_PAC_ strength and the behavioral improvement (*r* = 0.67, *p* = 4.9×10^-2^, Person’s correlation; Figure 5C), indicating that a stronger PPLE relates to a greater improvement in subjects’ performance. Collectively, PPLE exhibit a strong correlation with enhanced behavioral performance characterized by improved response stability and higher reaction speed.

## Discussion

Reward expectancy facilitates critical cognitive processes, such as attention, memory, and motivation^2,4,6^. To explore the neural basis of reward expectancy, we investigated iEEG activities in the AIC as subjects approached the pre-determined reward zones in a virtual T-maze task. Firstly, we identified pre-activation of RBPs in the AIC prior to reward onset. We further detected PAC between theta (5-7 Hz) oscillations and gamma (75–110 Hz) activity, with PAC strength exhibiting a positive correlation with RBP pre-activation levels. Notably, theta-gamma PAC emerged later than RBP pre-activation. Strikingly, peak gamma activities derived from PAC analysis progressively shifted to earlier theta phases across successive trials of approaching the same reward, exhibiting PPLEs. Furthermore, PPLE strength predicted trial-by-trial improvements in behavioral performance (e.g., response latency), highlighting its functional significance. These key findings uncover PPLE as a novel electrophysiological mechanism in the AIC, bridging temporal coordination of oscillatory activity with reward expectancy. The dynamic interplay between RBP pre-activation, theta-gamma PAC, and behavioral adaptation advances our understanding of how the AIC orchestrates reward expectancy (Figure 6).

**Figure 6.**
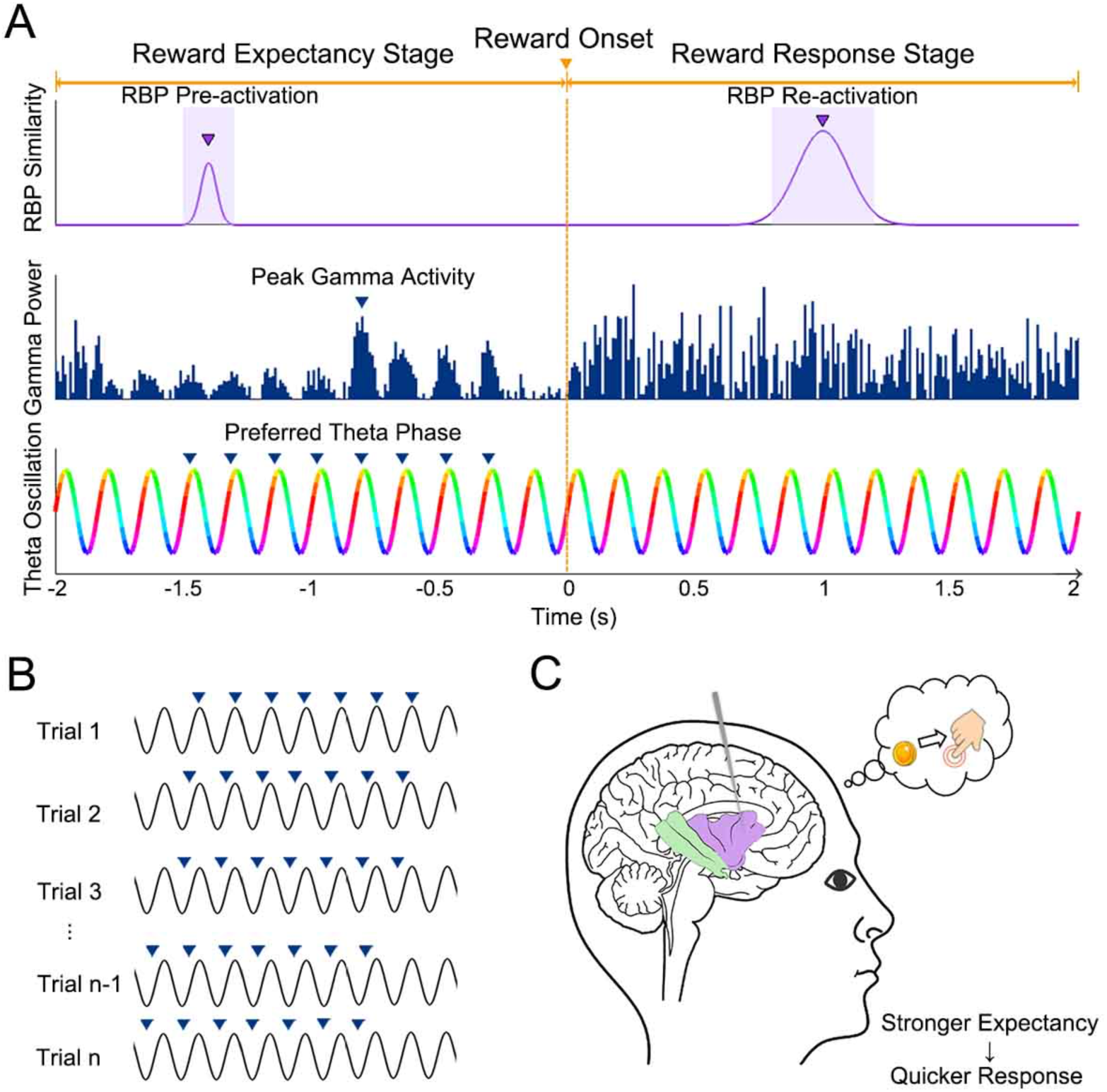
Neural-behavioral mechanisms of reward expectancy in the anterior insular cortex. (A) Single-trial neural dynamics. Upper panel: Reward-specific brain pattern (RBP, purple trace) shows anticipatory pre-activation (shaded area) during reward expectancy stage, peaking at the critical timepoint (purple triangle). Middle panel: Gamma-band power (blue bars) exhibits phase-dependent modulation by theta oscillations. The peak gamma activity (blue triangle) coupled to the preferred theta phase emerges temporally after RBP pre-activation. Bottom panel: Theta oscillations with continuous phase encoding (0–2π gradient color map). Blue triangles mark the preferred phase for gamma coupling. (B) Trial-to-trial phase-precession-like effect (PPLE). Individual trials demonstrate progressive phase advancement of theta-gamma coupling (blue triangles) across successive reward experiences. (C) Behavior-neural correlation. Subjects with stronger PPLE display faster behavioral responses to reward cues.

### Sequential Neural Activities during Reward Expectancy in the AIC

Recent neuroimaging evidence has demonstrated that the AIC is engaged during the early stages of reward processing^54–56^. An invasive investigation in non-human primates further identified a distinct class of AIC neurons that exhibit anticipatory firing patterns prior to the delivery of immediate rewards^16^. Complementing these findings, research in humans has revealed temporally structured neural activity within the AIC that underlies reward prediction^10^.

Our study revealed sequential pre-activation of RBPs during reward expectancy, with the AIC following hippocampal activation prior to subjects entering the reward zone (Figure 2C). This temporal hierarchy aligns with the SOCRATIC model, which posits that reward-related memories - conceptualized as RBP - are preactivated before repeated entering reward-associated contexts. Within this framework, theta-gamma PAC facilitates the integration of higher-order cognitive representations (e.g., reward expectancy) with lower-level sensory and contextual inputs, enabling temporal compression of real-world events into neural timescales^57–59^. This integration process underpins the formation of context-reward association^60^, and refines expectations related to rewards^3,14,61^. Consistent with this model, we observed enhanced coupling between theta (5-7 Hz) phase and gamma (75-110 Hz) power in the AIC, following the pre-activation of RBPs during reward expectancy stage (Figure 3A, B and E). Furthermore, the strength of PAC effect positively correlated with the pre-activation level of RBP (Figure 3C and D), suggesting that theta oscillations scaffold gamma-band synchronization to amplify reward-predictive signals. These findings support a hierarchical organization in which theta-gamma PAC and RBP pre-activation cooperatively mediate reward expectancy (Figure 6A), with theta rhythms priming gamma coordination for precise temporal encoding. The AIC’s role as a hub for integrating bottom-up contextual signals (e.g., spatial cues) with top-down prior knowledge (e.g., reward-related memory) is likely critical to this process^59^. Its extensive connectivity with limbic and neocortical regions positions it to dynamically enhance the salience of expected rewards and allocate cognitive resources (e.g., attention and motivation)^4,54^. Mechanistically, theta-gamma PAC may enable the AIC to “fast-forward” neural processing, generating predictive representations of rewards ahead of their actual occurrence^59^.

Theta-gamma coupling has been proposed modulated by dopaminergic pathways^58,59^. Recent neurophysiological studies in patients with Parkinson’s disease (PD) demonstrated that theta-gamma decoupling correlated with impaired reward processing, characterized by diminished reward positivity amplitudes and reduced striatal dopaminergic innervation^62^. Notably, dopaminergic medication partially restores theta-gamma coupling strength in PD, while tACS shows promise in modulating cross-frequency dynamics for cognitive remediation^62^. The AIC, which exhibits dense dopaminergic innervation, plays a critical role in reward prediction by integrating temporal and contextual cues to generate reward expectancy contains^63,64^. This aligns with its proposed function as a hub for translating bottom-up sensory inputs (e.g., spatial cues) into top-down anticipatory signals via theta-gamma PAC^59^. In contrast, the hippocampus prioritizes novel or unpredicted stimuli, a process driven by transient dopamine bursts that enhance theta-gamma coupling during exploratory behaviors^65,66^. This mechanistic distinction may explain our findings that hippocampal theta-gamma PAC peaks before entering decision zones (unpredicted stimulus; Figure 1C), whereas no significant PAC modulation occurs before entering reward zones (predetermined outcomes; Figure S2).

Collectively, these findings strengthen the evidence for the functional engagement of the AIC in anticipating forthcoming rewards, which is achieved through the establishment of context-reward association and saliency-based allocation of cognitive resources^4,60^. Our study provides the initial evidence of sequential and parallel coding for reward expectancy in the humans’ AIC. However, the causal relations of the sub-processes of reward expectancy remains un-addressed in the current study, which needs to be studied in further research.

### Phase Precession Effect of Macroscopic Brain Signals

The phase precession effect was initially identified in microscopic neural signaling, characterized by the progressive advancement of single-unit firing phases across successive theta cycles within sustained local field potentials^67,68^. This phenomenon has since been documented across diverse species, including rodents, non-human primates, and humans^39,42,46,69^. While most prominently observed in the medial temporal lobe, phase precession has also been detected in higher-order regions such as the prefrontal cortex and anterior cingulate cortex^32,41,46^. It has been posited as a critical mechanism for cognitive processing, facilitating the temporal compression of behavioral sequences into accelerated neural representations and enabling predictive coding of upcoming stimuli^41,42,46,67,68,70^. Inspiringly, recent findings indicate that the strength of phase precession correlates with the success of participants’ memory encoding and retrieval^46^.

Recent studies have challenged the traditional view of phase precession as a single-neuron property, reframing it as a manifestation of network-level coordination within distributed neural ensembles^71,72^. Unraveling the mechanisms and functional roles of macroscopic phase precession - particularly in large-scale cortical networks - remains critical for elucidating how neural dynamics govern cognition and behavior. High gamma activity (HGA), a proxy for population-level neuronal firing due to its tight coupling with spiking activity, offers a macroscopic lens to investigate such phenomena^73,74^. Nevertheless, direct evidence for theta phase precession effect of HGA remains sparse. A notable advance comes from Zheng et al., who identified cross-regional theta-gamma interactions during facial perception using a contextual modulation task^24^. Their work revealed a macroscopic phase precession effect, extending this phenomenon beyond single-unit spiking to population-level HGA and broadening its relevance to non-spatial cognitive states (e.g., transitions between task phases like context maintenance and face perception). In the present study, we identified analogous temporally structured HGA in the AIC during reward anticipation. Specifically, the peak gamma activities derived from PAC analysis were found systematically advanced to earlier theta phases as rewards approached across trials, termed PPLE (Figure 4A and 6B). This finding establishes PPLE as a multi-scale neural principle, spanning spatial domains (from single-unit spikes to HGA) and temporal scales (from millisecond oscillations to second-long behavioral epochs). Intriguingly, a recent neuromodulation study has demonstrated that the preferred phase for motor excitability progressively advances across successive tACS blocks^72^, providing independent empirical support for phase precession mechanisms observable at macroscopic scales. These observations provide a unifying framework for exploring how phase precession organizes neural activity across resolutions, with implications for understanding cognitive processes and neuro-modulatory interventions.

More exactly, PPLE in the AIC were related to reward expectancy, rather than reward outcome or a particular stimulus or location (Figure 4C and D), which further supports the involvement of the AIC in reward expectancy. In contrast, PPLE in the hippocampus did not exhibit any preference for reward expectancy (Figure 4C and D). This finding is analogous to the observation in rodents that the phase precession effect in the hippocampus is uniformly distributed and not affected by reward expectancy^42,43^. Nevertheless, the exact relationship between PPLE and phase precession effect remains unclear, due to lacking simultaneous recordings at micro- and macroscopic levels. Future studies combining macro- and micro-electrodes are valuable to determine whether PPLE have the same spatiotemporal distribution and functional role as theta phase precession effect.

### Reward Expectancy Promote Behavioral Performance

Reward expectancy has been suggested to significantly contribute to allocating cognitive resources and optimizing adaptive motor outputs^4,6,10,43^, enabling faster and more accurate behavioral responses to reward-related cues in individual subjects^2^. A prerequisite for the facilitation effect is the establishment the context-reward association through associative learning^2^. When subjects encounter a conditioned context, neural representations of the context are activated, triggering pre-activation of RBP and subsequent reward expectancy^3^. This process prioritizes attentional and motivational resources toward expected rewards, leveraging their salience within contextual frameworks to prime preparatory motor and cognitive states^3,4,54^. Consequently, subjects exhibit enhanced behavioral performance, marked by reduced reaction times and increased accuracy, due to preconfigured motor readiness^75^. Furthermore, reward outcomes that align with expectations reinforce existing context-reward associations through feedback mechanisms^2,60^.

In this study, the observed theta phase-gamma amplitude coupling and pre-activations of RBP during reward approach suggest a neural signature of context-reward association formation. These effects are proposed to conceptualize a temporal remapping of real-world events onto compressed neural timescales, thereby facilitating reward anticipation. We therefore assume that the presence and strength of PPLE reflect the degree of reward expectancy. To investigate their behavioral relevance, we conducted a preliminary analysis of PPLE in the AIC. Our results revealed that PPLE exerts a positive influence on behavioral performance (e.g., improved response speed; Figure 5C), underscoring the mechanistic role of reward expectancy in optimizing outcomes^1,2,4^. This finding highlights the potential for neuromodulation strategies to amplify PPLE, thereby augmenting cognitive and behavioral flexibility (Figure 6C).

To advance these findings, future research should prioritize multidimensional reward paradigms incorporating diverse reward types (e.g., psychological, material) and parametric variations in intensity, salience, and temporal predictability. Such frameworks will enable precise characterization of the spatiotemporal dynamics underlying reward expectancy, including oscillatory (e.g., PAC) and representational (e.g., RBP) mechanisms. This knowledge could inform the development of closed-loop neuromodulation protocols to selectively enhance context-reward associations and optimize behavioral performance in real-world scenarios.

### Conclusions

Our findings establish the AIC as a central hub orchestrating reward expectancy, integrating hierarchical neural mechanisms to bridge predictive coding with adaptive behavior. We reveal a temporally ordered cascade: RBP pre-activation primes the AIC, followed by theta-gamma PAC that amplifies reward-predictive signals. Critically, we identify PPLE - a novel electrophysiological signature defined by progressive shifts of peak gamma activates to earlier theta phases of the theta cycle across successive trials. This temporal recalibration mechanism enhances behavioral performance (e.g., shorten response latency), underscoring its role in optimizing reward expectancy.

## Methods

### Subject Details

Thirteen medically refractory epilepsy patients with depth electrodes for diagnosis (six males; average age = 25.25±8.87 years [mean ± SD]; Table S1) were recruited from the Hôpital de la Salpêtrière in Paris and the Second Affiliated Hospital of Zhejiang University in China. The study was approved by the Medical Ethics Committee at the hospital sites (INSERMC11-16 and StudyNo.2020-910, respectively), and written informed consent was obtained from all subjects.

### Experimental Design

We built a virtual T-maze (Panda3d and PandaEPL) in which subjects could freely navigate. As illustrated in Figure 1B, the T-maze consisted of 2 lateral roads, a central road, a starting zone, a decision zone, and three reward zones. The starting zone was both the beginning and the end of each trial. At the decision zone, subjects chose the correct direction to the lateral road according to the sign on the wall (circle or triangle represents left-turn or right-turn, respectively, Figure 1C and D). Among three reward zones, one was on the central road where a gold coin (value of 0.15 Euros) would definitely appear, and two were on the lateral roads where a gold coin appeared only if subjects chose the correct direction at the decision zone. Once the gold coin appeared, subjects were encouraged to press the Enter button to pick it up. Otherwise, the reward disappeared after subjects navigated through the reward zone. Subjects were encouraged to collect enough gold coins (10 Euros) as soon as possible. To avoid being disoriented, subjects were asked to memorize as many pictures hanging on the wall as possible to encourage them to pay attention to the task and their on-going position. Furthermore, different paintings of wallpapers also helped subjects to allocate themselves inside the maze to avoid being disoriented (as indicated in Figure 1D). We tested their knowledge about the location of pictures at the end of the experiment (data not reported here). Each subject required over an hour to complete the experiment.

During the experiment, subjects were positioned comfortably on their beds, with a laptop placed on a table directly in front of them. They used arrow buttons on the keyboard to navigate the virtual maze (Figure 1C). The full linear running speed was 6 virtual units/second and the linear acceleration speed was 40 virtual units/second^2^. The distance from the center of the starting zone to the center of the decision zone was 32 virtual units (Figure 1B, left). The distance between the center of the left arm and the center of the right arm measured 24 virtual units. The width of the corridor was 4 virtual units. Every time subjects picked up the reward or entered the decision zone, they paused for 2 seconds before continuing navigating. The left- and right-turn signs at the decision zone were counter-balanced within each subject. During the experiment, the testing laptop sent triggers to iEEG recording system in order to synchronize behavioral and iEEG data offline.

### Experimental Procedure

At the beginning of each trial, subjects stood at the starting location (Figure 1D). They first navigated through the middle arm of the T-maze toward the decision zone. On the middle arm, they encountered the first reward. Subsequently, subjects proceeded to enter the decision zone located at the other end of the middle corridor. Immediately after entering the decision zone, a triangle or a circle was signed on the wall. Subjects were asked to turn to the correct lateral arm in order to pick up the second reward of that trial. If they made a wrong choice at the decision zone (i.e., running into a wrong arm), the second reward of that trial would not appear. At the end of each trial, they navigated back to the starting zone for the next trial. Before the experiment, subjects performed a practice session by going through a different maze to familiarize themselves with the procedure of the experiment and navigating in the virtual environment.

### Behavioral Data Collection and Process

During the experiment, navigation trajectories of subjects were logged. We focused on three events as three experimental conditions for further behavioral and iEEG data analysis: entering the reward zone (ERZ), picking up the reward (PR) and entering the decision zone (EDZ). When subjects entered the reward zone, a gold coin appeared on the monitor. The event corresponding to PR was defined as the moment when subjects pressed the Enter button to collect rewards. Additionally, upon EDZ, a picture was displayed on the wall. To minimize the influence of learning biases, we specifically focused on the reward zone in the central corridor, where subjects’ expectations for receiving a reward were consistently fulfilled (i.e., the reward would certainly appear). Another advantage is ensuring the balance among the trial numbers under the three conditions included in the analysis. Nevertheless, ERZ events occurred on the lateral roads were also extracted to identify the reward-specific neural patterns (Figure 2A, see *Star Methods - Representational Similarity Analysis and Pre-activation of Reward-Specific Brain Patterns*). As a behavioral parameter, the response latency for picking up a reward was calculated for each ERZ event, referring to the time interval between the appearance of a gold coin and the button press to collect the coin in each trial (Figure 1F).

### Electrode Reconstruction

The AdTech electrodes used at the Hôpital de la Salpêtrière were equipped with 4-12 contacts, each measuring 1 mm in diameter, 2.41 mm in length, and spaced 5 mm apart^76^. Meanwhile, the HKHS electrodes employed at the Second Affiliated Hospital of Zhejiang University consisted of 8-16 contacts, with a diameter of 0.8 mm, length of 2 mm, and an inter-contact distance of 1.5 mm^77^.

Contacts located in the AIC and hippocampus, which are known to have close associations with reward processing and spatial memory^15,27,28^, were selected for further data analyses (Figure 1A). The exact location of each contact was defined via visual inspection in the local space of each subject using FSLeyes (https://fsl.fmrib.ox.ac.uk/fsl/fslwiki/FSLeyes).

After removing contacts at seizure onset zones and those contaminated by epileptic spikes, we obtained 25 AIC contacts (18 in the right hemisphere) from 9 subjects and 19 hippocampal contacts (10 in the right hemisphere) from 7 subjects (Figure 1A and Table S1). Three subjects (subject #8, #14 and #15) had both AIC and hippocampal contacts.

### IEEG Data Acquisition and Preprocessing

IEEG data was acquired using an Ad-Tech system (Hôpital de la Salpêtrière) and a Nihon-Kohden recording system (Second Affiliated Hospital of Zhejiang University) in a bipolar layout at sampling rates ζ1k Hz (Table S1). IEEG data were first down-sampled to 1kHz, bandpass filtered from 0.01 to 200 Hz, and notch filtered at 50 Hz and its harmonics. The filtered data were then re-referenced to the average activity of all depth electrodes outside seizure onset zones^47^. We segmented the iEEG data into epochs, spanning 2 seconds before to 2 seconds after each event of interest (i.e., ERZ, PR and EDZ). To remove any edge effects caused by time-frequency decomposition, one extra second before the start and after the end of each epoch was extracted. Data preprocessing was performed using EEGLAB toolbox and custom MATLAB code^78^.

Valid epochs were screened for each experimental condition. Epochs with breaks for clinical purposes (e.g., taking medicine; 1 epoch from subject #8 under ERZ condition), missing rewards (i.e., running out of the reward zone without picking up the reward; 1 epoch from subject #15 under PR condition), and incorrect responses at the decision zone (i.e., turning into a wrong arm; 4 epochs from subject #16 and 1 epoch from subject #17 under EDZ condition) were excluded. Epochs contaminated by artifacts such as epileptic activities were detected by visual inspection and excluded from further data analysis. The numbers of trials included were 32.08 ± 2.40 (mean ± SD) for ERZ, 32.23 ± 2.42 for PR, and 32.23 ± 2.39 for EDZ, while the numbers of trials excluded were 4.00 ± 2.86 for ERZ, 3.62 ± 2.69 for PR, and 3.46 ± 2.14 for EDZ.

### Representational Similarity Analysis and Pre-activation of Reward-Specific Brain Patterns

To illustrate the relevance of neural activities preceding reward appearance to reward expectancy, we employed representational similarity analysis (RSA) to test for the pre-activation of reward-specific brain pattern (RBP) prior to the reward onset (Figure 2A)^47–50^.

Initially, we identified RBP by analyzing post-ERZ events at three different spatial locations, including the mid-arm, left arm, and right arm (Figure 1B). To do this, we segmented each post-ERZ event into 200ms windows with 25ms overlap, yielding 73 time-windows per post-ERZ period. For each window, we measured the iEEG power across low-frequency bands (1-30 Hz in steps of 1 Hz) and high-frequency bands (35–150 Hz in steps of 5 Hz), creating a concatenated vector of 54 frequencies. This resulted in a 54 (frequency bins) ×73 (time bins) matrix for each post-ERZ event at each contact. We then computed the neural similarity between each post-ERZ/post-ERZ pair. This was done by computing between the frequency patterns of each time window in one trial (*m*) and each window in another trial (*n*) for each contact. The Fisher-z transformed correlation coefficients were averaged across all time windows for each *trial n*, resulting in a 1×73 (post-ERZ time bins) matrix for each post-ERZ/post-ERZ pair at each contact (RSA_ERZ/ERZ_; Figure 2A). To eliminate the effects of temporal proximity and location confounds, we excluded correlation coefficients derived from the same trials and those from the same location. Using the same approach, we also compared the neural patterns of post-ERZ events with post-EDZ events (RSA_ERZ/EDZ_) for each contact.

Subsequently, we conducted a multi-level linear regression analysis between RSA_ERZ/ERZ_ values and RSA_ERZ/EDZ_ values, accounting for variance at both the subject and contact levels for each post-ERZ time bin (Figure 2B). We considered bins with *p* < 0.05 as significant and grouped adjacent significant bins into clusters. The statistical value of each cluster was determined by summing the t-values from the regression analysis. To estimate the significance of these clusters, we employed a cluster-based permutation analysis^52^. This involved randomly shuffling the labels of post-ERZ or post-EDZ events for 500 times. For each permutation, we performed the same multi-level linear regression analysis and identified the cluster with the largest statistical value as the surrogate cluster. The significance level of the empirical clusters was then estimated by ranking their statistical value within the distribution of 500 surrogate clusters. An empirical cluster was considered significant if its statistical value was greater than the 95^th^ percentile of the surrogate distribution. The iEEG activities from time windows that exhibited significant higher RSA_ERZ/ERZ_ values than RSA_ERZ/EDZ_ values were identified as RBPs.

If reward information was processed before reward onset, we assumed that RBPs were pre-activated. In practice, we expected to observe that the pre-ERZ brain patterns were similar to RBPs, showing significant positive correlation coefficients. To determine the pre-activation of RBP during pre-ERZ periods, we computed the neural similarity between each pre-ERZ time bin’s brain patterns and RBP using Spearman’s correlation for each contact and trial. This procedure generated a time course of neural similarity values (RSA_preERZ/RBP_) over the pre-ERZ time windows. We then applied the same method to compare the neural patterns of pre-EDZ events with the RBP, yielding RSA_preEDZ/RBP_ values. To statically evaluate the difference between RSA_preERZ/RBP_ and RSA_preEDZ/RBP_, we conducted multi-level linear regression analysis considering subject and contact-level variances for each pre-ERZ time bin (Figure 2E). Bins with a p-value less than 0.05 were deemed significant and were grouped into clusters of adjacent significant bins. The statistical value of each cluster was calculated by summing the t-values from the regression analysis. To assess the significance of these clusters, we utilized a cluster-based permutation analysis, randomly re-assigning the labels of pre-ERZ or pre-EDZ events 500 times^52^. For each permutation, we repeated the multi-level linear regression and identified the cluster with the highest statistical value as a surrogate. The empirical clusters’ significance was determined by comparing their statistical values to the distribution of 500 surrogate clusters. A cluster was considered significant if its value exceeded the 95^th^ percentile of the surrogate distribution. The iEEG activities from time windows showing significantly higher pre-ERZ/RBP similarities compared to the baseline were identified as the presence of RBP pre-activation.

### Phase-Amplitude Coupling Analysis and Identification of Peak Gamma Activities

To estimate the strength of phase-amplitude coupling (PAC), we calculated the modulation index (MI) based on the Kullbeck–Leiber distance, a measure of the divergence between the probability distribution of high-frequency amplitudes and a uniform distribution^51,79^. We initially applied finite impulse response (FIR) filtering to iEEG data, yielding dynamics foreach low-frequency band (1-12 Hz in 1 Hz steps, 12 samples; frequency_low_) and each high-frequency band (30–150 Hz in 5 Hz steps, 25 samples; frequency_high_) for each ERZ trial. In accordance with the Heisenberg-Gabor limit, the low-frequency signals were filtered with a 1-Hz bandwidth, while the high-frequency signals were filtered with a 24-Hz bandwidth^80,81^. Each filtered low-frequency band was visually inspected to confirm its sinusoidal nature. Subsequently, the instantaneous phases of the low frequency signals and the amplitudes of the high frequency signals were extracted using the Hilbert transformation. The low-frequency phases were then binned into eighteen 20° intervals (−180° to 180°), and MI values were computed for each low-frequency phase and high-frequency amplitude pair. The obtained MI was further normalized by calculating the z-score of 200 surrogate datasets, generated by randomly shuffling the trial labels of high-frequency signals. For time windows before and after each ERZ event, the z-scored MI, computed for multiple frequencies of phase and amplitude, is represented in a comodulogram (Figure 3A).

To explore the differences in PAC strength before and after ERZ event, we conducted a comparison of the z-scored MI between the pre- and post-ERZ periods. This analysis employed multi-level linear regression analysis, which accounted for variance at both the subject and contact levels for each frequency_low_-frequency_high_ bin. Bins corresponding to a *p* level smaller than 0.05 were selected^82^. Adjacent bins were clustered together. The statistical value of each MI cluster was the summed t value from the multi-level linear regression analysis. The significance level of the empirical MI clusters was estimated using a cluster-based permutation analysis^52^. Within each permutation, we randomly shuffled condition labels (i.e., pre-or post-ERZ event) and performed the same multi-level linear regression analysis. This generated a set of surrogate MI clusters. The MI cluster with the largest statistical value was selected. The permutation procedure was repeated 500 times, generating 500 surrogate MI clusters. The significant level of each empirical MI cluster was estimated by ranking its statistical value within the distribution of 500 surrogate MI clusters. If the statistical value of an empirical cluster exceeded the 95^th^ percentile of the surrogate MI distribution, it was labeled as a significant cluster (*p* < 0.05, Figure 3A). We applied the same analysis to PR and EDZ conditions in contacts located at the AIC and hippocampus (Figure S2).

Within the frequency_low_-frequency_high_ cluster identified via previous PAC strength analysis, the instantaneous high-frequency amplitudes were averaged within each phase bins (eighteen 20° intervals)^83^. For each pre-ERZ event, the preferred PAC phase was defined as the low-frequency phase bin exhibiting the maximum high-frequency amplitude. This analysis was performed independently for each trial and recording contact. Subsequently, the peak gamma activity - corresponding to the highest amplitude coupled to the preferred PAC phase - was identified, and its temporal alignment was compared to the timing of RBP pre-activation (Figure 3E and S3).

### Phase-Precession-Like Effect of Peak Gamma Activities

To examine the underlying characteristics of trial-wise PAC, we investigated the phase-precession-like effect (PPLE) over trials within pre-ERZ event. To this end, we conducted a circular-linear correlation analysis for each contact^41^, treating the number of trials as a linear variable and the preferred PAC phase from each corresponding trial as a circular variable (Figure 4A and S4). The significance level of the coefficient value was estimated by performing a surrogate analysis. Within each permutation, we randomly shuffled trial labels and then computed the circular-linear correlation again^52^. This generated one surrogate coefficient value. The permutation was repeated 500 times and we obtained 500 surrogate coefficient values. If the circular-linear correlation coefficient derived from the empirical data exceeded the 95^th^ percentile of the surrogate coefficient distribution, the empirical correlation value was considered significant, indicating the existence of PPLE. We performed a control analysis by applying the same analysis to PR and EDZ events in AIC and hippocampal contacts (Figure S6 and S7).

## Resource Availability

### Lead contact

Further information and requests for resources should be directed to and will be fulfilled by the lead contact, Hui Zhang.

### Materials availability

This study did not generate new unique reagents.

### Data and code availability

The data are confidential medical records. The custom-written Matlab and Python code supporting the findings of this study will be made available upon publication.

## Acknowledgement

We are grateful to the patients for participating in our study. We thank the EEG technologists and nurses for assisting with data transfer and care of affected individuals during the EEG recording. This work was supported by the National Natural Science Foundation of China (NSFC; Grant Nos. 81901319, 81971207 and 82171437) and the Deutsche Forschungsgemeinschaft of Germany (DFG; Grant Nos. ZH822/1-1 and 316803389 - SFB 1280, projects F01).

## Author Contributions

Conceptualization, L.L.Y., S.W., and H.Z.; Methodology, H.Z., L.L.Y., S.C., and N.A.; Software, L.L.Y., H.Y.Y. and H.Z.; Formal Analysis, L.L.Y.; Investigation, L.L.Y. and K.L.; Resources, V.N. and S.W.; Visualization, L.L.Y. and X.F.Y.; Writing, L.L.Y. and H.Z.; Funding Acquisition, L.L.Y., S.W. and H.Z.; Supervision, H.Z. and S.W.; Project Administration, H.Z. and S.W.

## Declaration of Interests

The authors declare no competing interests.

## Notes

### Competing Interest Statement

The authors have declared no competing interest.

### Summary of Updates

1. Expanded data availability: Incorporated neurophysiological recordings and behavioral metrics from an additional participant with anterior insular cortex (AIC) electrode coverage. 2. Methodological refinements: a) Optimized parameters and permutation testing protocols of representational similarity analysis (RSA) to confirm the pre-activation of reward-specific brain patterns for both AIC and hippocampus, demonstrating enhanced statistical power b) Implemented robust phase-amplitude coupling (PAC) quantification using validated normalization procedures c) Introduced FOOOF spectral parameterization as a complementary approach to MODAL for slow oscillation detection (Supplementary Information) 3.Structural and textual improvements: a) Reorganized manuscript architecture to emphasize hypothesis-driven narrative progression b) Strengthened theoretical foundations in Introduction through expanded coverage of predictive coding frameworks c) Enhanced interpretation of cross-regional dynamics in Results/Discussion sections with comparative literature analysis 4.Supplemental material updates: a)Added validation analyses comparing two oscillation detection methodologies 5.Visual enhancements: a)Introduced schematic diagram elucidating AIC's proposed role in reward expectancy mechanisms (Figure 6) b)Updated all figures with improved statistical visualizations

## Reference

1. Pessiglione, M., Schmidt, L., Draganski, B., Kalisch, R., Lau, H., Dolan, R.J., and Frith, C.D. (2007). How the brain translates money into force: a neuroimaging study of subliminal motivation. Science 316, 904–906. 10.1126/science.1140459.

2. Rowe, J.B., Eckstein, D., Braver, T., and Owen, A.M. (2008). How does reward expectation influence cognition in the human brain? J Cogn Neurosci 20, 1980–1992. 10.1162/jocn.2008.20140.

3. Bein, O., Gasser, C., Amer, T., Maril, A., and Davachi, L. (2023). Predictions transform memories: How expected versus unexpected events are integrated or separated in memory. Neurosci Biobehav Rev 153, 105368. 10.1016/j.neubiorev.2023.105368.

4. Frömer, R., Lin, H., Dean Wolf, C.K., Inzlicht, M., and Shenhav, A. (2021). Expectations of reward and efficacy guide cognitive control allocation. Nat Commun 12, 1030. 10.1038/s41467-021-21315-z.

5. Blaukopf, C.L., and DiGirolamo, G.J. (2007). Reward, context, and human behaviour. ScientificWorldJournal 7, 626–640. 10.1100/tsw.2007.122.

6. Baruni, J.K., Lau, B., and Salzman, C.D. (2015). Reward expectation differentially modulates attentional behavior and activity in visual area V4. Nat Neurosci 18, 1656–1663. 10.1038/nn.4141.

7. Grill-Spector, K., Henson, R., and Martin, A. (2006). Repetition and the brain: neural models of stimulus-specific effects. Trends Cogn Sci 10, 14–23. 10.1016/j.tics.2005.11.006.

8. Bergmann, N., Koch, D., and Schubö, A. (2019). Reward expectation facilitates context learning and attentional guidance in visual search. J Vis 19, 10. 10.1167/19.3.10.

9. Seeley, W.W., Menon, V., Schatzberg, A.F., Keller, J., Glover, G.H., Kenna, H., Reiss, A.L., and Greicius, M.D. (2007). Dissociable intrinsic connectivity networks for salience processing and executive control. J Neurosci 27, 2349–2356. 10.1523/jneurosci.5587-06.2007.

10. Man, V., Cockburn, J., Flouty, O., Gander, P.E., Sawada, M., Kovach, C.K., Kawasaki, H., Oya, H., Howard III, M.A., and O’Doherty, J.P. (2024). Temporally organized representations of reward and risk in the human brain. Nature Communications 15, 2162.

11. Sridharan, D., Levitin, D.J., and Menon, V. (2008). A critical role for the right fronto-insular cortex in switching between central-executive and default-mode networks. Proc Natl Acad Sci U S A 105, 12569–12574. 10.1073/pnas.0800005105.

12. Cai, W., Chen, T., Ryali, S., Kochalka, J., Li, C.S., and Menon, V. (2016). Causal Interactions Within a Frontal-Cingulate-Parietal Network During Cognitive Control: Convergent Evidence from a Multisite-Multitask Investigation. Cereb Cortex 26, 2140–2153. 10.1093/cercor/bhv046.

13. Dajani, D.R., and Uddin, L.Q. (2015). Demystifying cognitive flexibility: Implications for clinical and developmental neuroscience. Trends Neurosci 38, 571–578. 10.1016/j.tins.2015.07.003.

14. Molnar-Szakacs, I., and Uddin, L.Q. (2022). Anterior insula as a gatekeeper of executive control. Neurosci Biobehav Rev 139, 104736. 10.1016/j.neubiorev.2022.104736.

15. Rothkirch, M., Schmack, K., Deserno, L., Darmohray, D., and Sterzer, P. (2014). Attentional modulation of reward processing in the human brain. Hum Brain Mapp 35, 3036–3051. 10.1002/hbm.22383.

16. Mizuhiki, T., Richmond, B.J., and Shidara, M. (2012). Encoding of reward expectation by monkey anterior insular neurons. J Neurophysiol 107, 2996–3007. 10.1152/jn.00282.2011.

17. Tort, A.B., Komorowski, R.W., Manns, J.R., Kopell, N.J., and Eichenbaum, H. (2009). Theta-gamma coupling increases during the learning of item-context associations. Proc Natl Acad Sci U S A 106, 20942–20947. 10.1073/pnas.0911331106.

18. Axmacher, N., Henseler, M.M., Jensen, O., Weinreich, I., Elger, C.E., and Fell, J. (2010). Cross-frequency coupling supports multi-item working memory in the human hippocampus. Proc Natl Acad Sci U S A 107, 3228–3233. 10.1073/pnas.0911531107.

19. Canolty, R.T., and Knight, R.T. (2010). The functional role of cross-frequency coupling. Trends Cogn Sci 14, 506–515. 10.1016/j.tics.2010.09.001.

20. McGinn, R.J., and Valiante, T.A. (2014). Phase-amplitude coupling and interlaminar synchrony are correlated in human neocortex. J Neurosci 34, 15923–15930. 10.1523/jneurosci.2771-14.2014.

21. Lisman, J. (2005). The theta/gamma discrete phase code occuring during the hippocampal phase precession may be a more general brain coding scheme. Hippocampus 15, 913–922. 10.1002/hipo.20121.

22. González, J., Cavelli, M., Mondino, A., Rubido, N., Bl Tort, A., and Torterolo, P. (2020). Communication Through Coherence by Means of Cross-frequency Coupling. Neuroscience 449, 157–164. 10.1016/j.neuroscience.2020.09.019.

23. Mariscal, M.G., Levin, A.R., Gabard-Durnam, L.J., Xie, W., Tager-Flusberg, H., and Nelson, C.A. (2021). EEG Phase-Amplitude Coupling Strength and Phase Preference: Association with Age over the First Three Years after Birth. eNeuro 8. 10.1523/eneuro.0264-20.2021.

24. Zheng, J., Skelin, I., and Lin, J.J. (2022). Neural computations underlying contextual processing in humans. Cell Rep 40, 111395. 10.1016/j.celrep.2022.111395.

25. Lopour, B.A., Tavassoli, A., Fried, I., and Ringach, D.L. (2013). Coding of information in the phase of local field potentials within human medial temporal lobe. Neuron 79, 594–606. 10.1016/j.neuron.2013.06.001.

26. Bush, D., and Burgess, N. (2020). Advantages and detection of phase coding in the absence of rhythmicity. Hippocampus 30, 745–762. 10.1002/hipo.23199.

27. Lopes-Dos-Santos, V., van de Ven, G.M., Morley, A., Trouche, S., Campo-Urriza, N., and Dupret, D. (2018). Parsing Hippocampal Theta Oscillations by Nested Spectral Components during Spatial Exploration and Memory-Guided Behavior. Neuron 100, 940–952.e947. 10.1016/j.neuron.2018.09.031.

28. Kragel, J.E., VanHaerents, S., Templer, J.W., Schuele, S., Rosenow, J.M., Nilakantan, A.S., and Bridge, D.J. (2020). Hippocampal theta coordinates memory processing during visual exploration. Elife 9. 10.7554/eLife.52108.

29. Rayan, A., Donoso, J.R., Mendez-Couz, M., Dolón, L., Cheng, S., and Manahan-Vaughan, D. (2022). Learning shifts the preferred theta phase of gamma oscillations in CA1. Hippocampus 32, 695–704. 10.1002/hipo.23460.

30. Siegel, M., Warden, M.R., and Miller, E.K. (2009). Phase-dependent neuronal coding of objects in short-term memory. Proc Natl Acad Sci U S A 106, 21341–21346. 10.1073/pnas.0908193106.

31. Yang, A.I., Dikecligil, G.N., Jiang, H., Das, S.R., Stein, J.M., Schuele, S.U., Rosenow, J.M., Davis, K.A., Lucas, T.H., and Gottfried, J.A. (2021). The what and when of olfactory working memory in humans. Curr Biol 31, 4499–4511.e4498. 10.1016/j.cub.2021.08.004.

32. Reddy, L., Self, M.W., Zoefel, B., Poncet, M., Possel, J.K., Peters, J.C., Baayen, J.C., Idema, S., VanRullen, R., and Roelfsema, P.R. (2021). Theta-phase dependent neuronal coding during sequence learning in human single neurons. Nat Commun 12, 4839. 10.1038/s41467-021-25150-0.

33. Watrous, A.J., Deuker, L., Fell, J., and Axmacher, N. (2015). Phase-amplitude coupling supports phase coding in human ECoG. Elife 4. 10.7554/eLife.07886.

34. Heusser, A.C., Poeppel, D., Ezzyat, Y., and Davachi, L. (2016). Episodic sequence memory is supported by a theta-gamma phase code. Nat Neurosci 19, 1374–1380. 10.1038/nn.4374.

35. Bahramisharif, A., Jensen, O., Jacobs, J., and Lisman, J. (2018). Serial representation of items during working memory maintenance at letter-selective cortical sites. PLoS Biol 16, e2003805. 10.1371/journal.pbio.2003805.

36. Freunberger, R., Werkle-Bergner, M., Griesmayr, B., Lindenberger, U., and Klimesch, W. (2011). Brain oscillatory correlates of working memory constraints. Brain Res 1375, 93–102. 10.1016/j.brainres.2010.12.048.

37. Lisman, J.E., and Idiart, M.A. (1995). Storage of 7 +/- 2 short-term memories in oscillatory subcycles. Science 267, 1512–1515. 10.1126/science.7878473.

38. Akkad, H., Dupont-Hadwen, J., Kane, E., Evans, C., Barrett, L., Frese, A., Tetkovic, I., Bestmann, S., and Stagg, C.J. (2021). Increasing human motor skill acquisition by driving theta-gamma coupling. Elife 10. 10.7554/eLife.67355.

39. Skaggs, W.E., McNaughton, B.L., Wilson, M.A., and Barnes, C.A. (1996). Theta phase precession in hippocampal neuronal populations and the compression of temporal sequences. Hippocampus 6, 149–172. 10.1002/(sici)1098-1063(1996)6:2<149::Aid-hipo6>3.0.Co;2-k.

40. Hafting, T., Fyhn, M., Bonnevie, T., Moser, M.B., and Moser, E.I. (2008). Hippocampus-independent phase precession in entorhinal grid cells. Nature 453, 1248–1252. 10.1038/nature06957.

41. Qasim, S.E., Fried, I., and Jacobs, J. (2021). Phase precession in the human hippocampus and entorhinal cortex. Cell 184, 3242–3255.e3210. 10.1016/j.cell.2021.04.017.

42. van der Meer, M.A., and Redish, A.D. (2011). Theta phase precession in rat ventral striatum links place and reward information. J Neurosci 31, 2843–2854. 10.1523/jneurosci.4869-10.2011.

43. Lansink, C.S., Meijer, G.T., Lankelma, J.V., Vinck, M.A., Jackson, J.C., and Pennartz, C.M. (2016). Reward Expectancy Strengthens CA1 Theta and Beta Band Synchronization and Hippocampal-Ventral Striatal Coupling. J Neurosci 36, 10598–10610. 10.1523/jneurosci.0682-16.2016.

44. Tingley, D., and Buzsáki, G. (2018). Transformation of a Spatial Map across the Hippocampal-Lateral Septal Circuit. Neuron 98, 1229–1242.e1225. 10.1016/j.neuron.2018.04.028.

45. Speers, L.J., Schmidt, R., and Bilkey, D.K. (2022). Aberrant Phase Precession of Lateral Septal Cells in a Maternal Immune Activation Model of Schizophrenia Risk May Disrupt the Integration of Location with Reward. J Neurosci 42, 4187–4201. 10.1523/jneurosci.0039-22.2022.

46. Zheng, J., Yebra, M., Schjetnan, A.G.P., Patel, K., Katz, C.N., Kyzar, M., Mosher, C.P., Kalia, S.K., Chung, J.M., and Reed, C.M. (2024). Theta phase precession supports memory formation and retrieval of naturalistic experience in humans. Nature human behaviour. 10.1038/s41562-024-01983-9.

47. Zhang, H., Fell, J., and Axmacher, N. (2018). Electrophysiological mechanisms of human memory consolidation. Nat Commun 9, 4103. 10.1038/s41467-018-06553-y.

48. Kriegeskorte, N., Mur, M., and Bandettini, P. (2008). Representational similarity analysis - connecting the branches of systems neuroscience. Front Syst Neurosci 2, 4. 10.3389/neuro.06.004.2008.

49. Xue, G., Dong, Q., Chen, C., Lu, Z., Mumford, J.A., and Poldrack, R.A. (2010). Greater neural pattern similarity across repetitions is associated with better memory. Science 330, 97–101. 10.1126/science.1193125.

50. Pacheco-Estefan, D., Fellner, M.-C., Kunz, L., Zhang, H., Reinacher, P., Roy, C., Brandt, A., Schulze-Bonhage, A., Yang, L., and Wang, S. (2024). Maintenance and transformation of representational formats during working memory prioritization. Nature Communications 15, 8234.

51. Tort, A.B., Komorowski, R., Eichenbaum, H., and Kopell, N. (2010). Measuring phase-amplitude coupling between neuronal oscillations of different frequencies. J Neurophysiol 104, 1195–1210. 10.1152/jn.00106.2010.

52. Zhang, H. (2023). How Can I Conduct Surrogate Analyses, and How Should I Shuffle? In Intracranial EEG: A Guide for Cognitive Neuroscientists, N. Axmacher, ed. (Springer International Publishing), pp. 567-577. 10.1007/978-3-031-20910-9_35.

53. Hermoso-Mendizabal, A., Hyafil, A., Rueda-Orozco, P.E., Jaramillo, S., Robbe, D., and de la Rocha, J. (2020). Response outcomes gate the impact of expectations on perceptual decisions. Nat Commun 11, 1057. 10.1038/s41467-020-14824-w.

54. Schneider, M., Leuchs, L., Czisch, M., Sämann, P.G., and Spoormaker, V.I. (2018). Disentangling reward anticipation with simultaneous pupillometry / fMRI. Neuroimage 178, 11–22. 10.1016/j.neuroimage.2018.04.078.

55. Apaydın, N., Üstün, S., Kale, E.H., Çelikağ, İ., Özgüven, H.D., Baskak, B., and Çiçek, M. (2018). Neural Mechanisms Underlying Time Perception and Reward Anticipation. Front Hum Neurosci 12, 115. 10.3389/fnhum.2018.00115.

56. O’Doherty, J.P., Deichmann, R., Critchley, H.D., and Dolan, R.J. (2002). Neural responses during anticipation of a primary taste reward. Neuron 33, 815–826. 10.1016/s0896-6273(02)00603-7.

57. Mariscal, M.G., Berry-Kravis, E., Buxbaum, J.D., Ethridge, L.E., Filip-Dhima, R., Foss-Feig, J.H., Kolevzon, A., Modi, M.E., Mosconi, M.W., Nelson, C.A., et al. (2021). Shifted phase of EEG cross-frequency coupling in individuals with Phelan-McDermid syndrome. Mol Autism 12, 29. 10.1186/s13229-020-00411-9.

58. John, E., Lisman, Nonna, A., and Otmakhova (2001). Storage, recall, and novelty detection of sequences by the hippocampus: Elaborating on the SOCRATIC model to account for normal and aberrant effects of dopamine. Hippocampus.

59. Köster, M. (2024). The theta-gamma code in predictive processing and mnemonic updating. Neurosci Biobehav Rev 158, 105529. 10.1016/j.neubiorev.2023.105529.

60. Sun, K., Xiao, L., Wu, Y., Zuo, D., Zhang, C., Liu, S., He, Z., Rong, S., Wang, F., and Sun, T. (2020). GABAergic neurons in the insular cortex play an important role in cue-morphine reward memory reconsolidation. Life Sci 254, 117655. 10.1016/j.lfs.2020.117655.

61. Uddin, L.Q., Kinnison, J., Pessoa, L., and Anderson, M.L. (2014). Beyond the tripartite cognition-emotion-interoception model of the human insular cortex. J Cogn Neurosci 26, 16–27. 10.1162/jocn_a_00462.

62. Sharma, R., and Thirugnanasambandam, N. (2025). Theta-Gamma Decoupling - A neurophysiological marker of impaired reward processing in Parkinson’s disease. Brain Res 1850, 149406. 10.1016/j.brainres.2024.149406.

63. Hurd, Y.L., Suzuki, M., and Sedvall, G.C. (2001). D1 and D2 dopamine receptor mRNA expression in whole hemisphere sections of the human brain. J Chem Neuroanat 22, 127–137. 10.1016/s0891-0618(01)00122-3.

64. Allman, J.M., Watson, K.K., Tetreault, N.A., and Hakeem, A.Y. (2005). Intuition and autism: a possible role for Von Economo neurons. Trends Cogn Sci 9, 367–373. 10.1016/j.tics.2005.06.008.

65. Knight, R.T. (1996). Contribution of human hippocampal region to novelty detection. Nature 383, 256–259. 10.1038/383256a0.

66. Mirenowicz, J., and Schultz, W. (1994). Importance of unpredictability for reward responses in primate dopamine neurons. Journal of Neurophysiology 72, 1024–1027.

67. O’Keefe, J., and Recce, M.L. (1993). Phase relationship between hippocampal place units and the EEG theta rhythm. Hippocampus 3, 317–330. 10.1002/hipo.450030307.

68. Jones, M.W., and Wilson, M.A. (2005). Phase precession of medial prefrontal cortical activity relative to the hippocampal theta rhythm. Hippocampus 15, 867–873. 10.1002/hipo.20119.

69. Courellis, H.S., Nummela, S.U., Metke, M., Diehl, G.W., and Miller, C.T. (2019). Spatial encoding in primate hippocampus during free navigation. PLoS Biology 17, e3000546.

70. Reifenstein, E.T., Bin Khalid, I., and Kempter, R. (2021). Synaptic learning rules for sequence learning. Elife 10. 10.7554/eLife.67171.

71. Weinrich, C.A., Brittain, J.S., Nowak, M., Salimi-Khorshidi, R., Brown, P., and Stagg, C.J. (2017). Modulation of Long-Range Connectivity Patterns via Frequency-Specific Stimulation of Human Cortex. Current Biology 27, 3061–3068.

72. Wischnewski, M., Tran, H., Zhao, Z., Shirinpour, S., Haigh, Z.J., Rotteveel, J., Perera, N.D., Alekseichuk, I., Zimmermann, J., and Opitz, A. (2024). Induced neural phase precession through exogenous electric fields. Nature Communications 15.

73. Ray, S., and Maunsell, J.H. (2011). Different origins of gamma rhythm and high-gamma activity in macaque visual cortex. PLoS Biol 9, e1000610. 10.1371/journal.pbio.1000610.

74. Belitski, A., Gretton, A., Magri, C., Murayama, Y., Montemurro, M.A., Logothetis, N.K., and Panzeri, S. (2008). Low-frequency local field potentials and spikes in primary visual cortex convey independent visual information. J Neurosci 28, 5696–5709. 10.1523/jneurosci.0009-08.2008.

75. Kok, P., Failing, M.F., and de Lange, F.P. (2014). Prior expectations evoke stimulus templates in the primary visual cortex. J Cogn Neurosci 26, 1546–1554. 10.1162/jocn_a_00562.

76. Lehongre, K., Lambrecq, V., Whitmarsh, S., Frazzini, V., Cousyn, L., Soleil, D., Fernandez-Vidal, S., Mathon, B., Houot, M., Lemaréchal, J.D., et al. (2022). Long-term deep intracerebral microelectrode recordings in patients with drug-resistant epilepsy: Proposed guidelines based on 10-year experience. Neuroimage 254, 119116. 10.1016/j.neuroimage.2022.119116.

77. Chen, C., Wang, Y., Ye, L., Xu, J., Ming, W., Liu, X., Hu, L., Ye, H., Xu, C., Wang, Y., et al. (2023). A region-specific modulation of sleep slow waves on interictal epilepsy markers in focal epilepsy. Epilepsia 64, 973–985. 10.1111/epi.17518.

78. Delorme, A., and Makeig, S. (2004). EEGLAB: an open source toolbox for analysis of single-trial EEG dynamics including independent component analysis. J Neurosci Methods 134, 9–21. 10.1016/j.jneumeth.2003.10.009.

79. Yin, Z., Zhu, G., Liu, Y., Zhao, B., Liu, D., Bai, Y., Zhang, Q., Shi, L., Feng, T., Yang, A., et al. (2022). Cortical phase-amplitude coupling is key to the occurrence and treatment of freezing of gait. Brain 145, 2407–2421. 10.1093/brain/awac121.

80. Aru, J., Aru, J., Priesemann, V., Wibral, M., Lana, L., Pipa, G., Singer, W., and Vicente, R. (2015). Untangling cross-frequency coupling in neuroscience. Curr Opin Neurobiol 31, 51–61. 10.1016/j.conb.2014.08.002.

81. Seymour, R.A., Rippon, G., and Kessler, K. (2017). The Detection of Phase Amplitude Coupling during Sensory Processing. Front Neurosci 11, 487. 10.3389/fnins.2017.00487.

82. Zhang, H., Fell, J., Staresina, B.P., Weber, B., Elger, C.E., and Axmacher, N. (2015). Gamma power reductions accompany stimulus-specific representations of dynamic events. Curr Biol 25, 635–640. 10.1016/j.cub.2015.01.011.

83. Port, R.G., Berman, J.I., Liu, S., Featherstone, R.E., Roberts, T.P.L., and Siegel, S.J. (2019). Parvalbumin Cell Ablation of NMDA-R1 Leads to Altered Phase, But Not Amplitude, of Gamma-Band Cross-Frequency Coupling. Brain Connect 9, 263–272. 10.1089/brain.2018.0639.

