## Supplementary Information for "Phase-precession-like Effect in the Anterior Insula Cortex during Reward Expectancy"

**The file includes:**

Spectral Dynamics of Theta and Alpha Oscillations in the Anterior Insular Cortex and Hippocampus

Tables S1. Demographic information.

Figures S1 to S8

References

### Spectral Dynamics of Theta and Alpha Oscillations in the Anterior Insular Cortex and Hippocampus

Before performing further analyses on the oscillatory data, we first explored the existence of low-frequency oscillations (LFOs) across experimental events (e.g., entering the reward zone [ERZ], picking up the reward [PR], and entering the decision zone [EDZ]) and recording contacts in the anterior insular cortex (AIC) and hippocampus (Figure S1). To this end, we performed the Multiple oscillations detection algorithm (MODAL) to confirm the presence of LFOs^1^. Firstly, power spectral density (1-50 Hz in steps of 0.5 Hz) was calculated using Morlet wavelet transformation (6 cycles) for each trial within each contact (Figure S1A). Then, a 1/f regression to the background spectrum was constructed in log-log space, and narrow-band oscillations exceeding the fitted line were identified^1^ (Figure S1B). To evaluate the prevalence of oscillations at each frequency, the percentage of detected time was calculated for each contact and trial which were further averaged across contacts and trials for each experimental condition and subject (Figure S1C-E). In the AIC, LFOs remained relatively constant and pronounced within event-centered time windows (Figure S1D). Theta oscillations (3.5-7.5Hz) were most prevalent, detected in nearly half of the time bins on average across subjects (ERZ: 41.39 ± 1.36% [mean ± SD], PR: 42.84 ± 1.62%, EDZ: 43.20 ± 2.24%). Less prevalent LFOs were observed in the alpha (8-12Hz; ERZ: 13.00 ± 1.25%, PR: 12.83 ± 1.24%, EDZ: 12.96 ± 1.81%) and beta (12.5-30 Hz; ERZ: 13.44 ± 1.55%, PR: 12.37 ± 1.08%, EDZ: 13.45 ± 1.73%) bands. The prevalence of theta oscillation significantly exceeded that alpha and beta oscillations, respectively for each experimental condition (Friedman test: *ps* ≤ 3×10^-4^; post-hoc Wilcoxon matched-pairs signed rank test: *ps* ≤ 3.9×10^-3^ for theta vs. alpha and theta vs. beta). In contrast, alpha and beta oscillations showed no significant difference in prevalence (post-hoc Wilcoxon matched-pairs signed rank test: *ps* ≥ 0.50).

Similar patterns emerged in the hippocampus during event-locked time windows (Figure S1E), with LFOs predominantly observed in the theta band (ERZ: 41.86 ± 0.96%, PR: 42.33 ± 1.06%, EDZ: 41.30 ± 1.04%), followed by alpha (ERZ: 13.46 ± 1.61%, PR: 12.78 ± 1.67%, EDZ: 13.36 ± 1.74%) and beta (ERZ: 13.02 ± 2.43%, PR: 42.33 ± 1.06%, EDZ: 12.68 ± 1.65%) bands. Theta oscillations exhibited significantly higher prevalence than alpha and beta oscillations across all experimental conditions (Friedman test: *ps* ≤ 1.2×10^-4^; post-hoc Wilcoxon matched-pairs signed rank test: *ps* ≤ 0.016 for theta vs. alpha and theta vs. beta). In contrast, no significant difference in prevalence was observed between alpha and beta oscillations (post-hoc Wilcoxon matched-pairs signed rank test: *ps* ≥ 0.16).

To further validate these findings, we performed the Fitting Oscillations & One Over F (FOOOF) algorithm^2^. For each trial and contact, power spectral density (PSD) was computed using Welch's method with Hamming windows, spaning 1-40 Hz in 0.5 Hz increments. The PSD with was decomposed into aperiodic and oscillatory constituents with the following parameters: aperiodic_mode = ‘fixed’, peak_width_limits = [1, 6], max_n_peaks = 6, peak_threshold = 2, and proximity_threshold = 1.5. Subtracting the aperiodic fit from the raw PSD revealed periodic peaks in the residual "flattened" spectrum (Figure S1F). Only peaks with an r-squared value greater than 0.95 and peak power higher than 0.6 a.u. were retained. The frequency distribution of identified peaks was quantified across experimental conditions in the AIC and hippocampus (Figure S1G and H). In the AIC, spectral peaks were predominantly clustered within the theta band across all conditions. In contrast, the hippocampus exhibited a broader spectral profile, with peaks spanning both theta and alpha bands, dynamically modulated by behavioral context.

Taken together, these findings reinforce the role of theta and alpha oscillations in cognition and align with prior reports of LFOs coordinating neural activity during virtual navigation^1,3,4^.

### Reference

1.	Watrous, A.J., Miller, J., Qasim, S.E., Fried, I., and Jacobs, J. (2018). Phase-tuned neuronal firing encodes human contextual representations for navigational goals. Elife *7*. 10.7554/eLife.32554.

2.	Donoghue, T., Haller, M., Peterson, E.J., Varma, P., Sebastian, P., Gao, R., Noto, T., Lara, A.H., Wallis, J.D., Knight, R.T., et al. (2020). Parameterizing neural power spectra into periodic and aperiodic components. Nat Neurosci *23*, 1655-1665. 10.1038/s41593-020-00744-x.

3.	Kahana, M.J., Sekuler, R., Caplan, J.B., Kirschen, M., and Madsen, J.R. (1999). Human theta oscillations exhibit task dependence during virtual maze navigation. Nature *399*, 781-784. 10.1038/21645.

4.	Bohbot, V.D., Copara, M.S., Gotman, J., and Ekstrom, A.D. (2017). Low-frequency theta oscillations in the human hippocampus during real-world and virtual navigation. Nat Commun *8*, 14415. 10.1038/ncomms14415.

### Table S1. Demographic information. Related to Figure 1

| **Subject number** | **Gender** | **Age** | **Handedness** | **Epileptic zone** | **Total number of electrodes** | **Sampling rate (Hz)** | **Number Electrodes included** | | **Trial numbers (included/excluded)** | | | **Sample source** |
| --- | --- | --- | --- | --- | --- | --- | --- | --- | --- | --- | --- | --- |
|  |  |  |  |  |  |  | **AIC** | **Hippocampus** | **ERZ** | **PR** | **EDZ** |  |
| 1 | f | 45 | Uncertain | Uncertain | 74 | 2048 | × | 5R2L | 33/6 | 32/7 | 35/2 | Hôpital de la Salpêtrière |
| 2 | f | 19 | Right | Medial temporal lobe (L) | 34 | 1000 | × | 2L | 31/3 | 30/4 | 30/4 | Hôpital de la Salpêtrière |
| 4 | f | 21 | Right | Cingulate Gyrus (R) | 58 | 4000 | 2R | × | 31/0 | 31/0 | 29/3 | Hôpital de la Salpêtrière |
| 7 | m | 37 | Right | Temporal lobe (R) | 50 | 4000 | 1R | × | 30/5 | 30/5 | 31/4 | Hôpital de la Salpêtrière |
| `8 | f | 18 | Right | Temporal Lobe (L) | 41 | 4000 | 1L | 1L | 29/6 | 30/5 | 29/5 | Hôpital de la Salpêtrière |
| 10 | m | 19 | Right | Temporal Lobe (L) | 65 | 4000 | × | 2L | 37/2 | 38/1 | 35/3 | Hôpital de la Salpêtrière |
| 12 | m | 29 | Left | Uncertain | 47 | 4000 | 4L | × | 30/6 | 31/5 | 32/4 | Hôpital de la Salpêtrière |
| 13 | m | 26 | Right | Temporal Lobe (R) | 66 | 1000 | × | 1L | 30/5 | 30/5 | 30/5 | Hôpital de la Salpêtrière |
| 14 | m | 22 | Right | Frontal/Temporal Lobe (R) | 64 | 1000 | 4R | 3R | 34/0 | 34/0 | 34/0 | Hôpital de la Salpêtrière |
| 15 | f | 29 | Left | Medial temporal lobe (L) | 82 | 1000 | 3R | 1R2L | 34/2 | 34/1 | 35/1 | Hôpital de la Salpêtrière |
| 16 | m | 25 | Right | Frontal lobe (R) | 53 | 2000 | 6R1L | × | 30/7 | 31/5 | 31/5 | Hôpital de la Salpêtrière |
| 17 | f | 13 | Right | Precentral sulci (R) | 86 | 2000 | 2R | × | 34/9 | 34/8 | 35/8 | Second Affiliated Hospital of Zhejiang University |
| 18 | f | 27 | Right | temporal lobe (L) | 54 | 4096 | 1L | × | 34/1 | 34/1 | 33/1 | Hôpital de la Salpêtrière |

Abbreviations: AIC, anterior insular cortex; ERZ, entering the reward zone; PR, picking up the reward; EDZ, entering the decision zone; R, right; L, left; f, female; m, male.

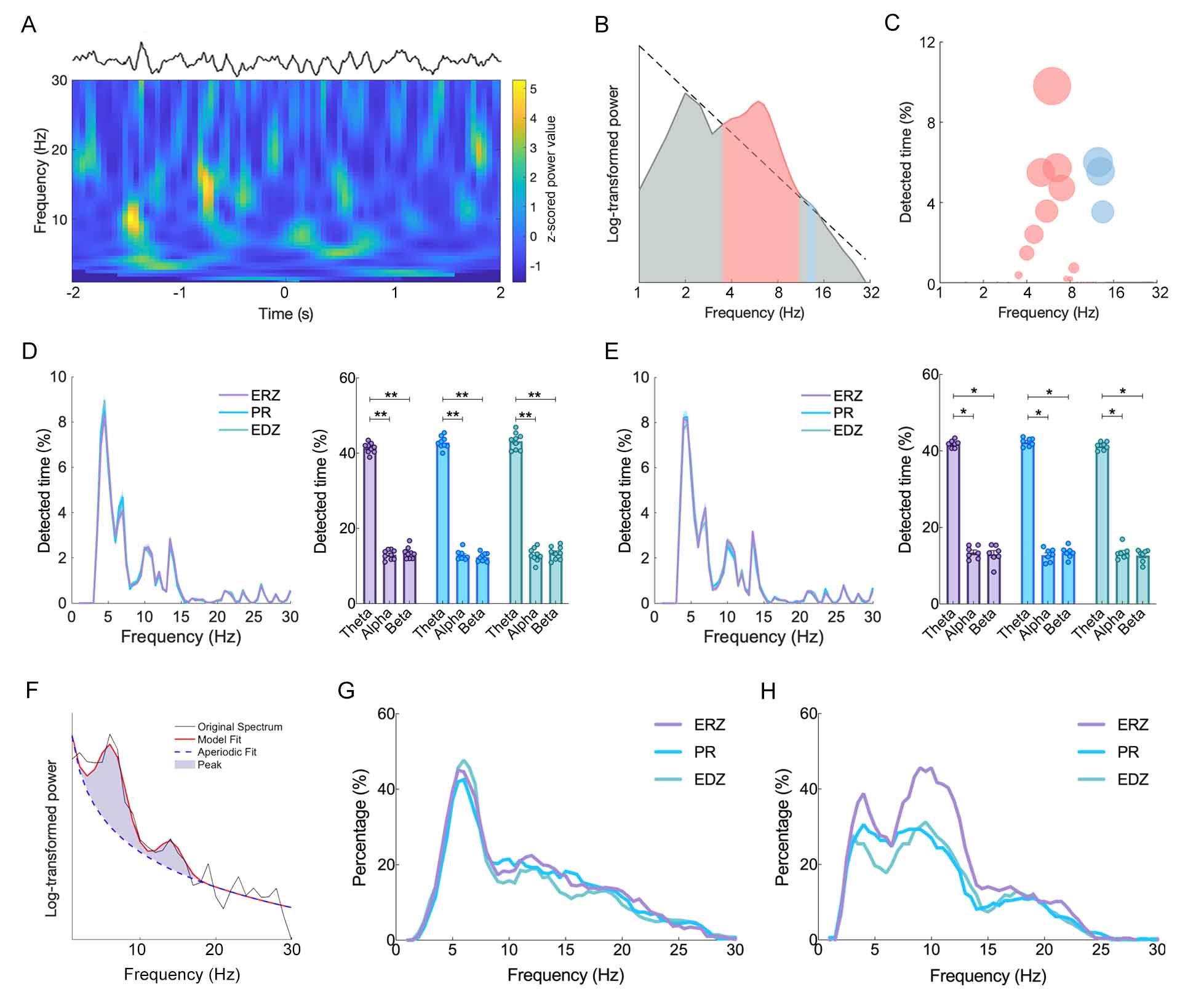

### Figure S1. Low-frequency oscillations detected in the anterior insular cortex and hippocampus during virtual navigation.

(A-C) An example trial from a recording contact in the right anterior insular cortex (AIC) of subject #16. (A) Raw trace and its time-frequency spectrum in the example trial. Zero represents the timepoint of entering the reward zone. (B) Two narrow-band oscillations (3.5-11Hz indicated by pink; 12.5-14Hz indicated by light blue) are identified as contiguous frequencies exceeding the fitted line (dashed line) using the Multiple oscillations detection algorithm (MODAL). (C) Prevalence of oscillations at each frequency from the trial shown in panel B. The larger the sphere, the higher the percentage of detected time of the corresponding frequency. (D-E) Population-level oscillation prevalence in AIC (D) and hippocampal (E) contacts. Left: Percentage of time periods during which oscillatory activities are detected separately for each frequency. Right: Proportion of time periods for each detected oscillation band (theta: 3.5-7.5Hz; alpha: 8-12Hz; beta: 12.5-30Hz). Rings indicate individual subject, and error bars indicate one standard deviation (SD) from the mean. *, 0.01≤*p*<0.05; **, 0.005≤*p*<0.01. (F) Example spectral decomposition using the Fitting Oscillations & One Over F (FOOOF) model. Narrow-band oscillations (light purple) are identified in the residual flattened spectrum after subtracting the aperiodic component (dashed line). (G-H) Frequency distribution of spectral peaks across experimental conditions in the AIC (G) and hippocampus (H). Each color denotes one experimental condition. ERZ, enter the reward zone; PR, picking up the reward; EDZ, enter the decision zone.

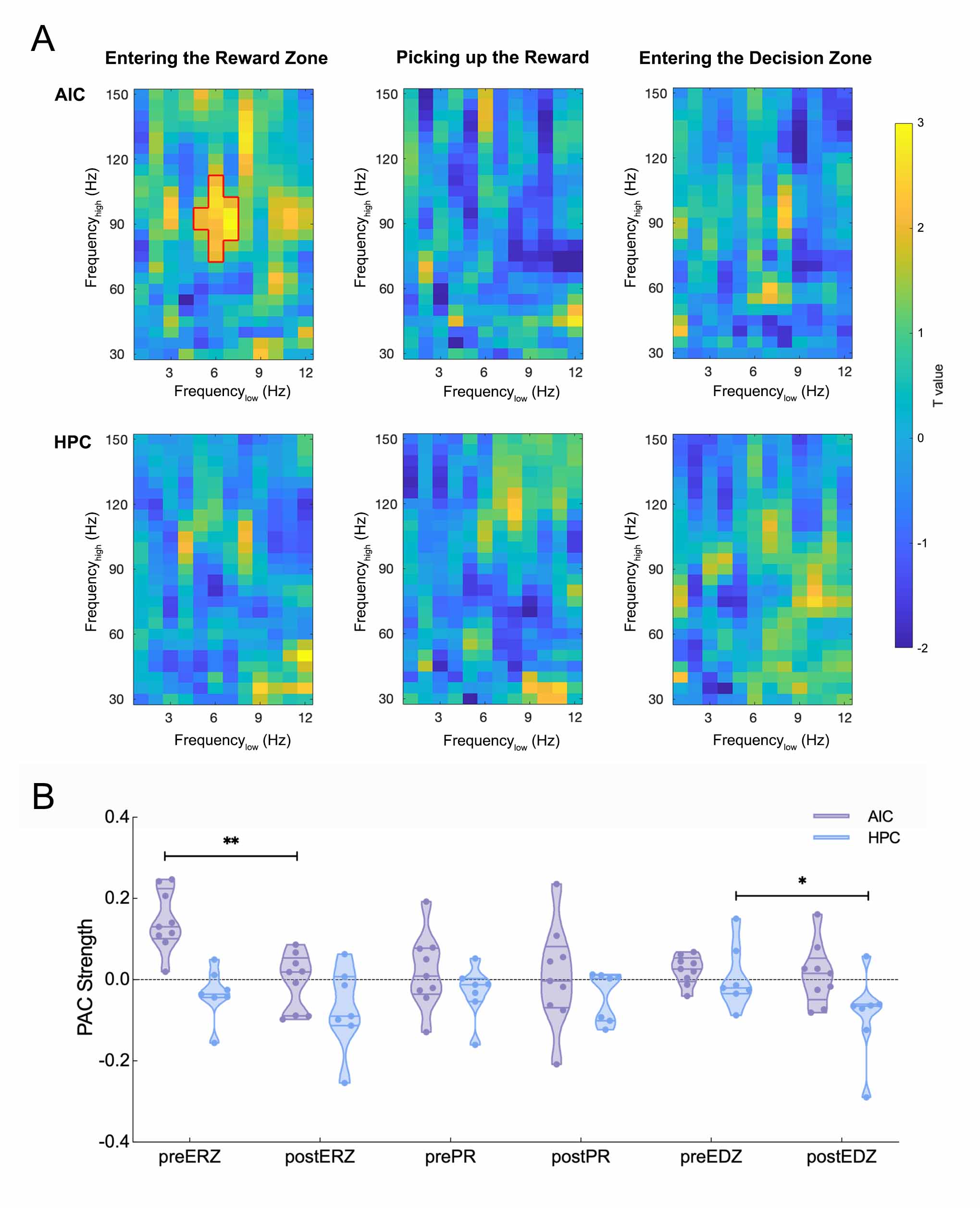

### Figure S2. PAC analysis of each experimental condition for contacts in the AIC and hippocampus separately. Related to Figure 3.

(A) Comodulograms showing the difference of PAC values between time windows before and after each event - entering the reward zone (ERZ; left), picking up the reward (PR; middle), entering the decision zone (EDZ; right). These are depicted separately for the anterior insular cortex (AIC, upper) and hippocampus (HPC, lower). Frequency_low_-frequency_high_ bins circled by red lines indicate enhanced PAC effects for the period before ERZ event in the AIC. (B) Average PAC values within the frequency_low_-frequency_high_ cluster from Figure 3A in the AIC (purple) and hippocampus (blue) during different experimental conditions. Each violin plot indicates one condition, and each dot within violin plots indicates one subject. *:0.01≤p<0.05; **: 0.005≤p<0.01.

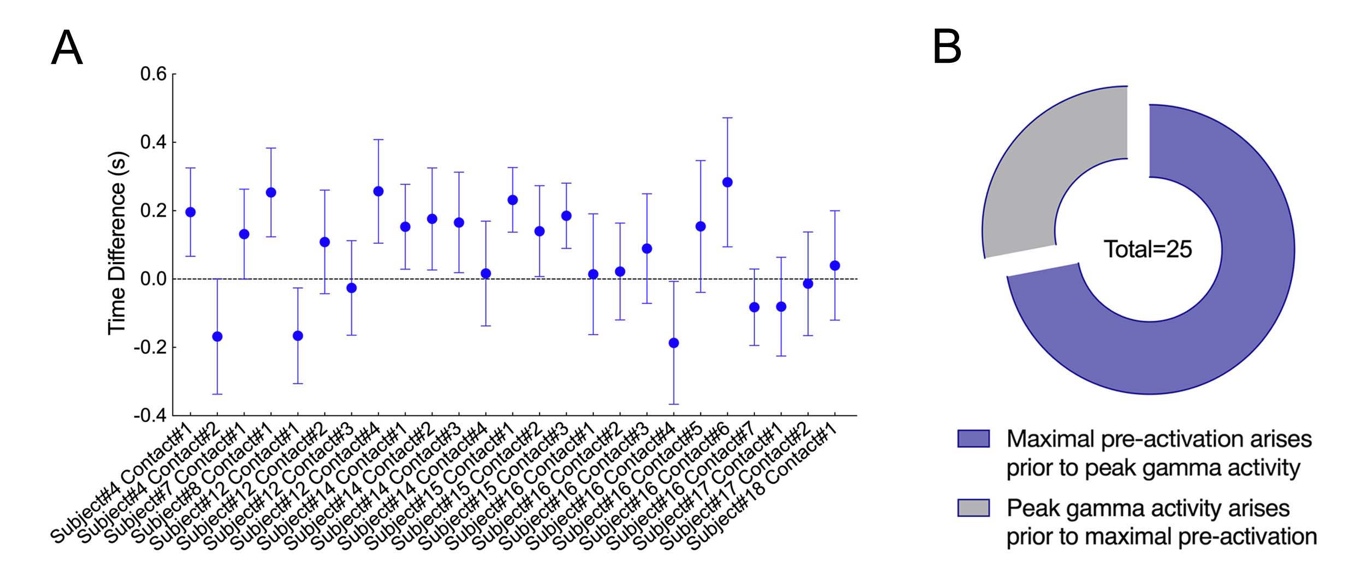

### Figure S3. Temporal relationship between PAC effect and RBP pre-activation. Related to Figure 3.

(A) Time difference between the peaked gamma activities locked to preferred PAC phase and the maximal pre-activation of reward-specific brain patterns (RBPs). Dots indicate individual recording contacts; error bars denote SEMs. (B) RBP pre-activation precedes gamma amplitude peaks in 72% (18/25, purple) of contacts.

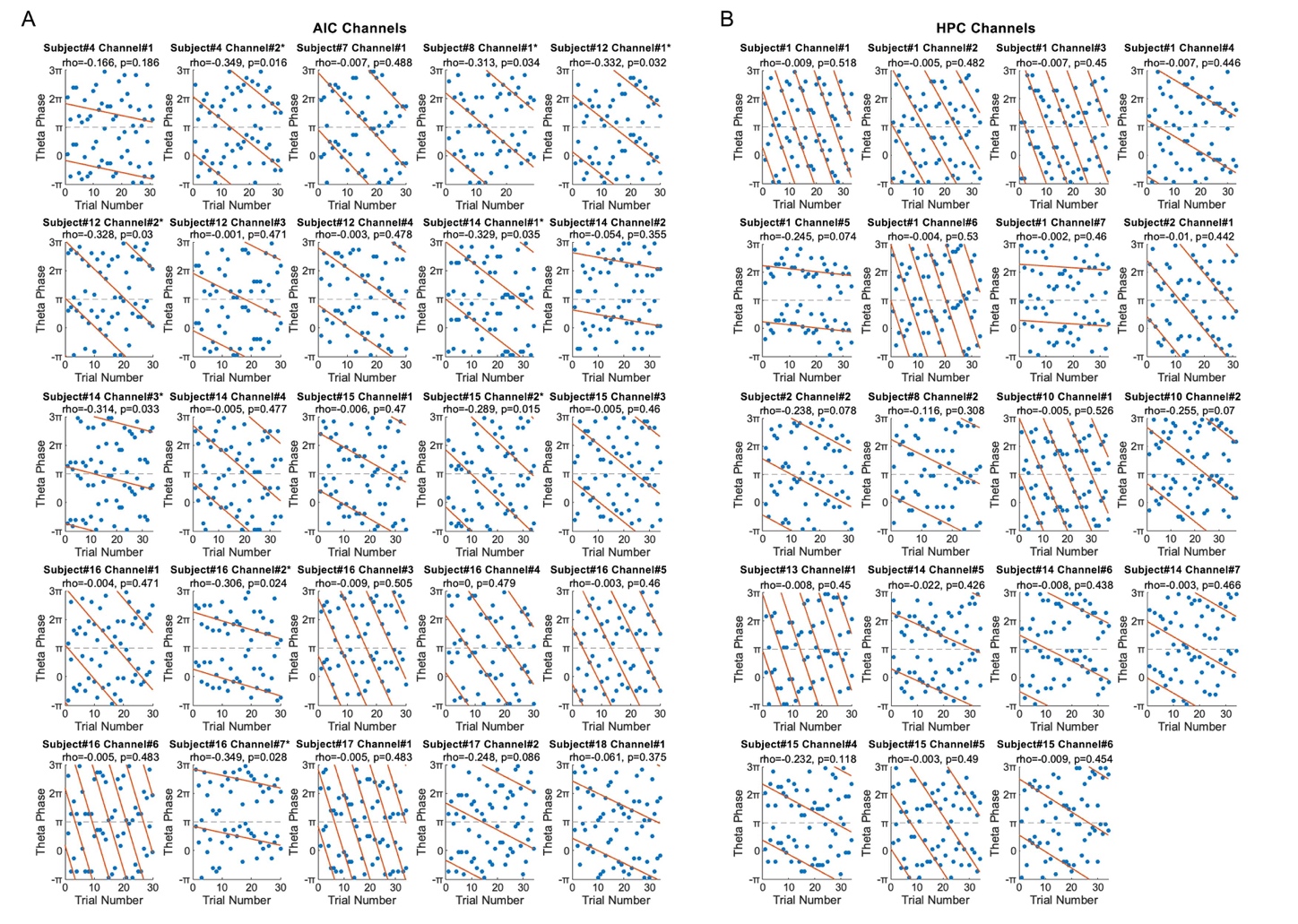

### Figure S4. PPLE during the period before entering reward zones in the anterior insular cortex and hippocampus. Related to Figure 4.

Phase-precession-like effect (PPLE) for the theta phase extracted from phase-amplitude coupling (PAC) analysis before entering reward zones for each contact in the anterior insular cortex (AIC; A) and hippocampus (HPC; B). Each panel displays data for each contact, with blue dots indicating individual trials and orange lines representing circular-linear regression fits. Asterisks denote significant effects.

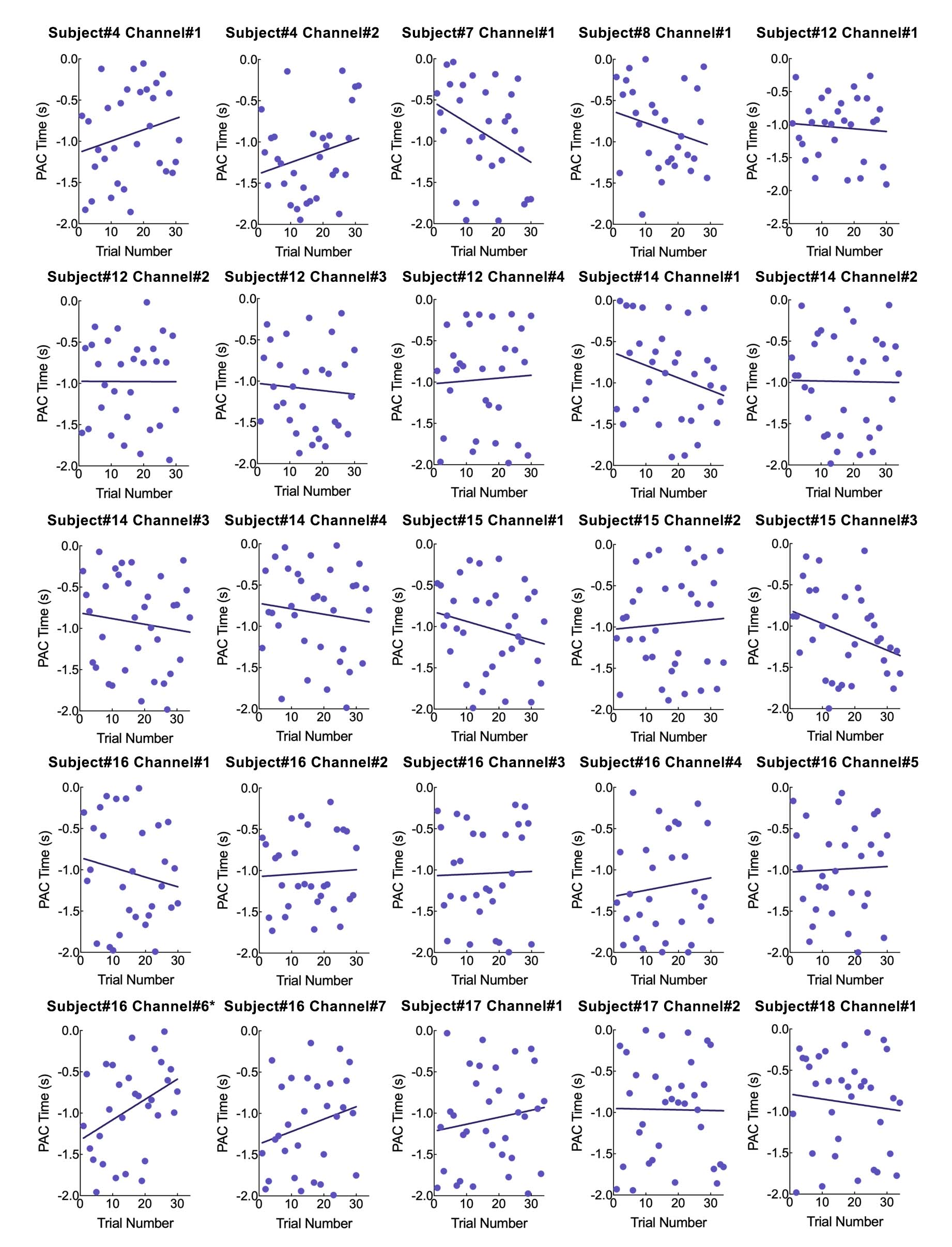

### Figure S5. Correlation between the time of peak gamma power and the number of trials for individual contact in the anterior insular cortex (AIC).

Each panel displays data for each contact, with blue dots indicating individual trials and blue lines representing simple linear regression fits. Y axis indicates the time course relative to pre-ERZ period, with time 0 marking the onset of rewards. Asterisks denote significant effects.

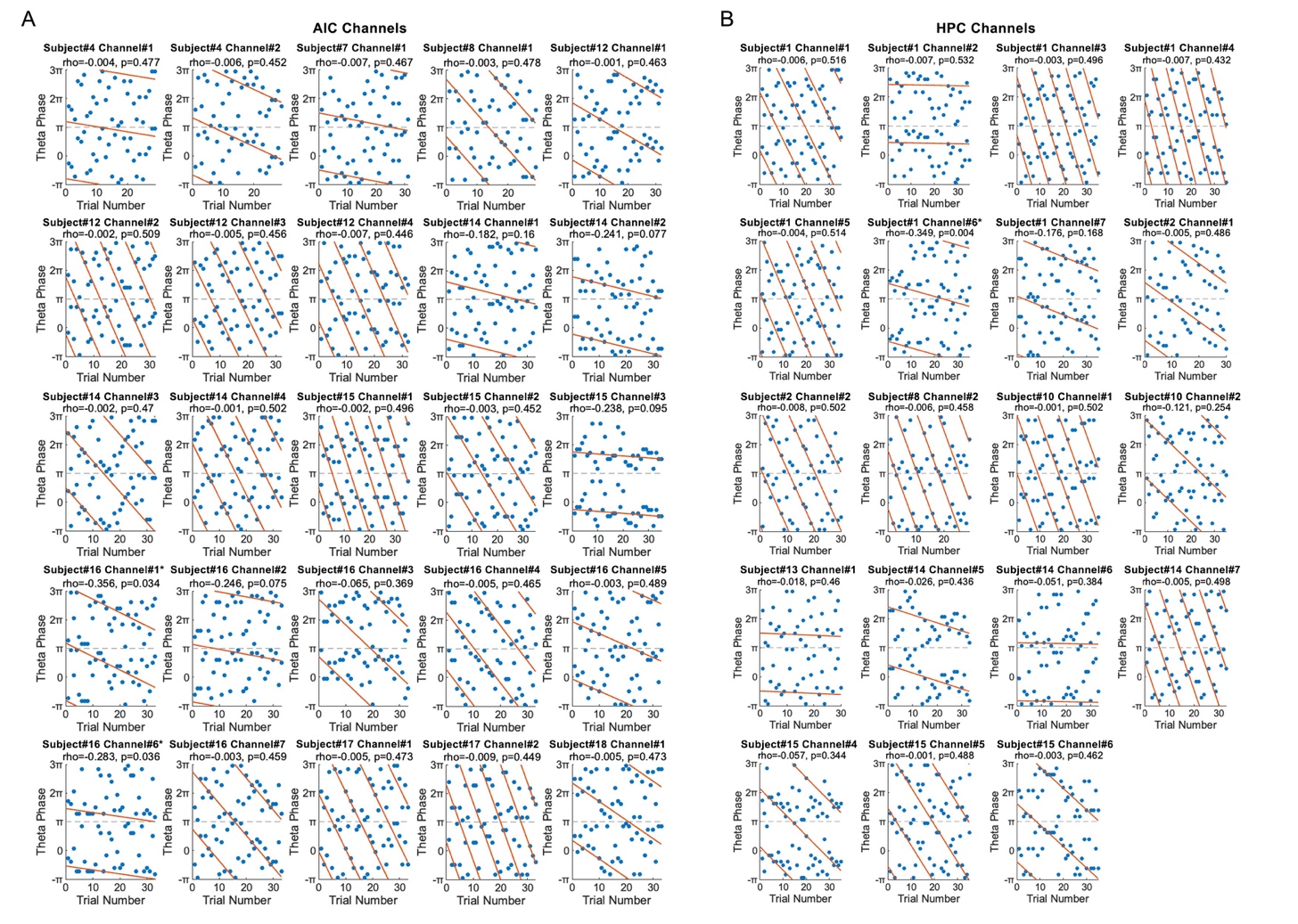

### Figure S6. PPLE during the period before entering decision zones in the anterior insular cortex and hippocampus. Related to Figure 4.

Phase-precession-like effect (PPLE) for the theta phase extracted from phase-amplitude coupling (PAC) analysis before entering decision zones for each contact in the anterior insular cortex (AIC; A) and hippocampus (HPC; B). Each panel displays data for each contact, with blue dots indicating individual trials and orange lines representing circular-linear regression fits. Asterisks denote significant effects.

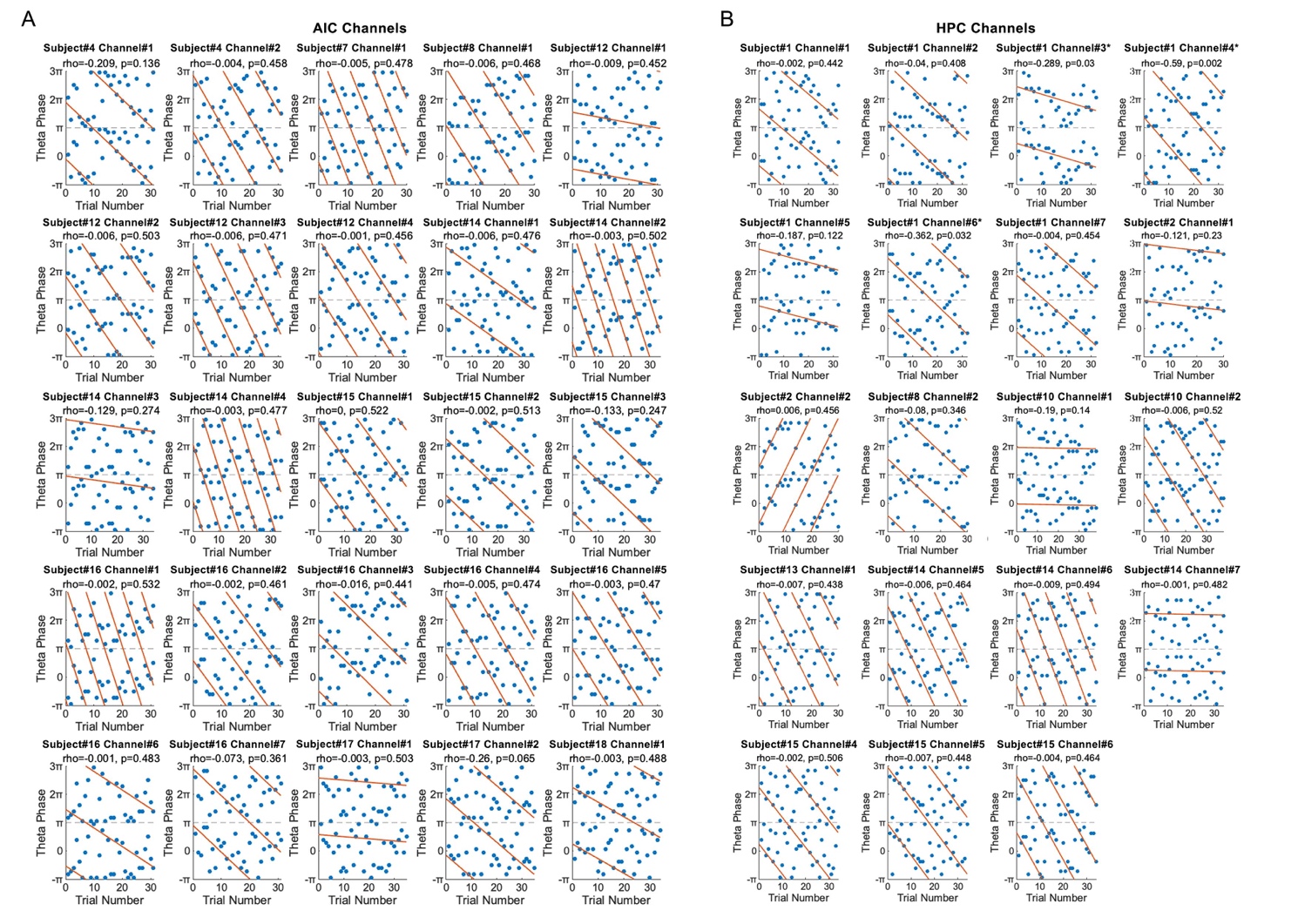

### Figure S7. PPLE during the period before picking up rewards in the anterior insular cortex and hippocampus. Related to Figure 4.

Phase-precession-like effect (PPLE) for the theta phase extracted from phase-amplitude coupling (PAC) analysis before picking up rewards for each contact in the anterior insular cortex (AIC; A) and hippocampus (HPC; B). Each panel displays data for each contact, with blue dots indicating individual trials and orange lines representing circular-linear regression fits. Asterisks denote significant effects.

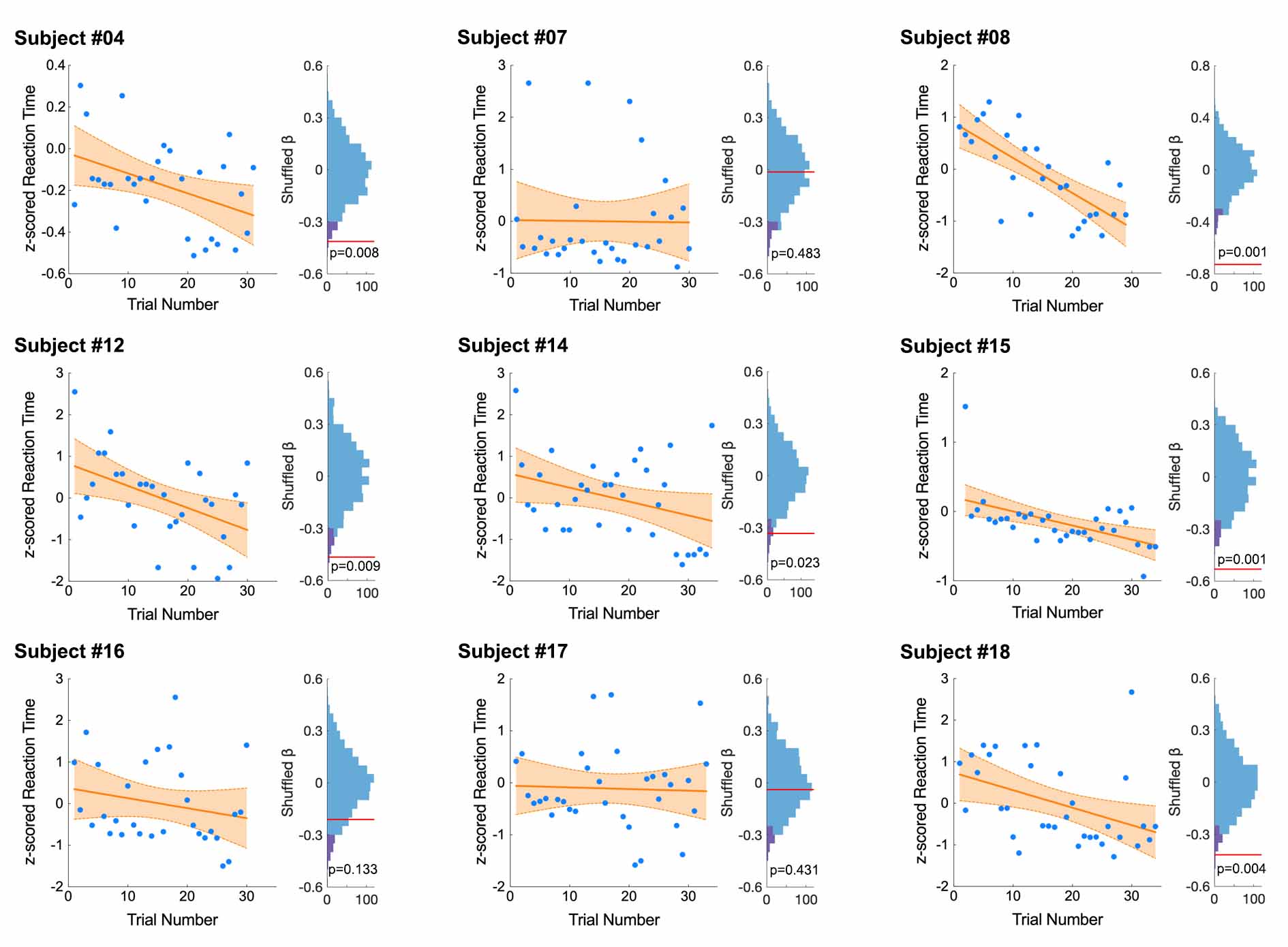

### Figure S8. Declining response time trends with trial numbers in subjects with anterior insular cortex contacts. Related to Figure 1 and 5.

The declining pattern of response times as a function of trial numbers in nine subjects with contacts in the anterior insular cortex. Each scatter-plot illustrates the relationship between response times and trial numbers for an individual subject, with each dot representing one trial. Orange line indicates the fitted regression line between response latency and trial numbers, and shaded area represents the 95% confidence interval of the fitted lines. To the right of each scatter plot, a bar-plot displays the rank of the empirical data (indicated by the red vertical line) within the surrogate distribution for each subject. Purple shaded area indicates the significance level.
